# Lineage-specific CDK activity dynamics characterize early mammalian development

**DOI:** 10.1101/2024.06.14.599098

**Authors:** Bechara Saykali, Andy D. Tran, James A. Cornwell, Matthew A. Caldwell, Paniz Rezvan Sangsari, Nicole Y. Morgan, Michael J. Kruhlak, Steven D. Cappell, Sergio Ruiz

## Abstract

Cyclin-dependent kinases (CDK) are key regulatory enzymes that regulate proliferation dynamics and cell fate in response to extracellular inputs. It remains largely unknown how CDK activity fluctuates and influences cell commitment *in vivo* during early mammalian development. Here, we generated a transgenic mouse model expressing a CDK kinase translocation reporter (KTR) that enabled quantification of CDK activity in live single cells. By examining pre- and post-implantation mouse embryos at different stages, we observed a progressive decrease in CDK activity in cells from the trophectoderm (TE) prior to implantation. This drop correlated with the establishment of an FGF4-dependent signaling gradient through the embryonic-abembryonic axis. Furthermore, we showed that CDK activity levels do not determine cell fate decisions during pre-implantation development. Finally, we uncovered the existence of conserved regulatory mechanisms in mammals by revealing lineage-specific regulation of CDK activity in TE-like human cells.

## INTRODUCTION

The cell cycle is tightly controlled by protein complexes of regulatory subunits, cyclins, which bind their catalytic partners, cyclin-dependent kinases (CDK) (Malumbres and Barbacid, 2009). In a classical model for cell cycle progression, stimulation with mitogens triggers the upregulation of D-type cyclins in G1 phase of the cell cycle, which activate CDK4 and CDK6 (Massague, 2004). Together with E- and A-type cyclins and their associated CDK (mainly CDK2 but also CDK1), cyclin D-CDK4/6 promote the phosphorylation and inactivation of the retinoblastoma protein RB1 leading to E2F-mediated transactivation of genes required to progress through S-phase (Fisher et al, 2022). Finally, at the end of the G2-phase, B-type cyclins translocate to the nucleus and activate CDK1 to promote entry into mitosis. An additional layer of control over CDK activity is imposed by the members of the INK4 (p16^Ink4a^, p15^Ink4b^, p18^Ink4c^ and p19^Ink4d^) and CIP/KIP (p27^kip1^, p21^cip1^ and p57^kip2^) families of CDK inhibitors which inhibit CDK4/6 or CDK1/2 complexes, respectively (Nakayama and Nakayama, 1998). In summary, cell cycle progression is precisely controlled by oscillatory CDK activity regulated by the timely expression of cyclins and CDK inhibitors.

Embryonic development is a prime example of precise control over cell division dynamics to regulate cell number and fate (Ciemerych and Sicinski, 2005). In approximately seventy-two hours, the mouse fertilized egg develops into a blastocyst, a self-organized hollow sphere with an outer polarized epithelium (trophectoderm, TE) enclosing a group of cells known as the inner cell mass (ICM) (Yao et al, 2019). Before implantation, the ICM further differentiates into primitive endoderm (PrE) and the epiblast (EPI). The TE and PrE will give rise to the extraembryonic cell types whereas the EPI will give rise to the entire fetus (Chazaud and Yamanaka, 2016; Rossant, 2018). At the 8-cell (8C) stage, cell compaction is the mechanical signal that induces the first lineage decision based on the mutually exclusive expression of CDX2 and SOX2, which specify TE and ICM, respectively. The second lineage decision occurs within the ICM at embryonic day 3.5 (E3.5) and involves the expression of NANOG, to specify the EPI, and GATA6 to specify the PrE (Rossant, 2018; Chazaud and Yamanaka, 2016). Although the duration of the first divisions following fertilization is comparable to that observed in somatic cells (around twenty hours), pronounced differences are observed at the blastocyst stage where EPI cells quickly divide approximately every ten hours (Ciemerych and Sicinski, 2005). Moreover, cells from the polar TE (those in close contact with the ICM) proliferate more rapidly than the cells from the mural TE (cells encompassing the blastocyst cavity) (Copp, 1978). After implantation, cells in the mural TE terminally differentiate into trophoblast giant cells (TGC), mononuclear but polyploid cells essential for embryo development. Cells in the polar TE remain multipotent and highly proliferative generating the extra-embryonic ectoderm (ExE) while EPI cells generate the egg cylinder with cells showing extremely short doubling times (five to eight hours), due to dramatic shortening of the G1 and G2 phases. Following gastrulation, the cell cycle length increases due to the extension of the G1 and G2 phases (Ciemerych and Sicinski, 2005).

Pre-implantation development is a robust and autonomous process independent of exogenous mitogens suggesting that regulatory pathways controlling the cell cycle may work differently in embryonic cells than in somatic cells (Stewart and Cullinan, 1997). Indeed, RB1 is undetectable from the 4C-stage until the late blastocyst (Iwamori et al, 2002), and the expression of CDK inhibitors is extremely low or non-existent before implantation (Ciemerych and Sicinski, 2005; Boroviak et al, 2015). These observations explain the fast kinetics of cell divisions and the shortened G1 and G2 phases during early development. Most of our knowledge on cell cycle regulation in the pre-implantation embryo was derived from studies on embryonic stem cells (ESC), the *in vitro* counterpart of the ICM (White and Dalton, 2005; Padgett and Santos, 2020). ESC proliferate at unusually rapid rates and have a characteristic cell cycle structure with truncated G1 and G2 phases recapitulating what has been observed in the ICM (Liu et al, 2019; White et al, 2005). They divide every ten to twelve hours and are characterized by high levels of cyclins E/A and CDK1/2 activity (Stead et al, 2002). Remarkably, cyclin E/A-CDK2 activity is constitutive throughout the entire cell cycle and lacks any periodicity in its regulation. Moreover, although the cyclin B-CDK1 complex retains some periodicity in its activity, it is substantially higher compared to the activity detected in somatic cells (Stead et al, 2002), and needed to maintain their epigenetic identity and pluripotent potential (Michowski et al, 2020). However, it is unclear whether embryonic cells from different lineages show oscillatory or constitutive CDK regulation *in vivo* or whether changes in CDK activity influence cell fate decisions during early lineage allocation in the embryo. Thus, we generated a mouse model containing a CDK sensor that enabled quantification of CDK activity via live imaging in embryos. While EPI cells retain elevated CDK activity, we observed a drastic reduction of CDK activity due to low FGF signaling in cells from the TE in blastocysts prior to implantation. Furthermore, we confirmed the lineage-specific downregulation of CDK activity in a system that recapitulates human gastrulation *in vitro*, demonstrating the existence of conservation in this process.

## RESULTS

### Generation of a mouse model with a live sensor to examine CDK activity

We used a previously described CDK sensor (Spencer et al, 2013) which includes a portion of the DNA human helicase B (DHB) containing four CDK phosphorylation sites fused to the fluorescent protein mClover3. We generated a transgene encoding for both the CDK sensor and the histone 2B (H2B) fused to the fluorescent protein mRuby3 separated from each other by a T2A self-cleaving peptide sequence to facilitate segmentation and tracking in time-lapse experiments (DHB/H2B hereafter, Figure 1A). This CDK sensor enabled live imaging and quantification of CDK activity according to its subcellular localization. As the cell progresses during the cell cycle, the fluorescent-based reporter translocates between the nucleus (low activity) and the cytoplasm (high activity) in a CDK-dependent manner, providing a quantifiable readout of CDK activity at a single-cell level (Figure 1A). We first developed, using CRISPR/Cas9 targeting, murine ESC carrying a constitutive transgene integrated at the *Rosa26* locus expressing our modified CDK sensor (ESC^DHB/H2B^; Figure 1A). Imaging the CDK sensor in proliferating ESC^DHB/H2B^ revealed that most cells showed cytoplasmic localization and confirmed the elevated levels of CDK activity driving the cell cycle in pluripotent cells (Figure 1B). To determine the levels of CDK activity, we used the cytoplasmic-to-nuclear intensity ratio (C/N) of the DHB-mClover3 chimeric protein as a proxy (Spencer et al, 2013). When we tracked individual ESC^DHB/H2B^ over time we observed that following mitosis, cells quickly recovered elevated levels of CDK activity and increased nuclear size prior to a new mitosis (Figure 1C). This observation is in accordance with the observed constitutive CDK activity throughout the entire cell cycle in ESC. To validate the responsiveness of the CDK sensor in pluripotent cells, ESC^DHB/H2B^ were treated with a CDK1/2 inhibitor (CDK1/2i) which promoted efficient CDK inhibition and nuclear accumulation of the reporter (Figure 1B and S1A). This accumulation was time and dose-dependent demonstrating the excellent dynamic range of the CDK sensor in ESC (Figure 1D). Furthermore, we also observed nuclear translocation when cells were treated with inhibitors for mTOR or MYC (Scognamiglio et al, 2016; Bulut-Karslioglu et al, 2016), which promote a quiescent-like state in ESC (Figure S1B and S1C).

**Figure 1:**
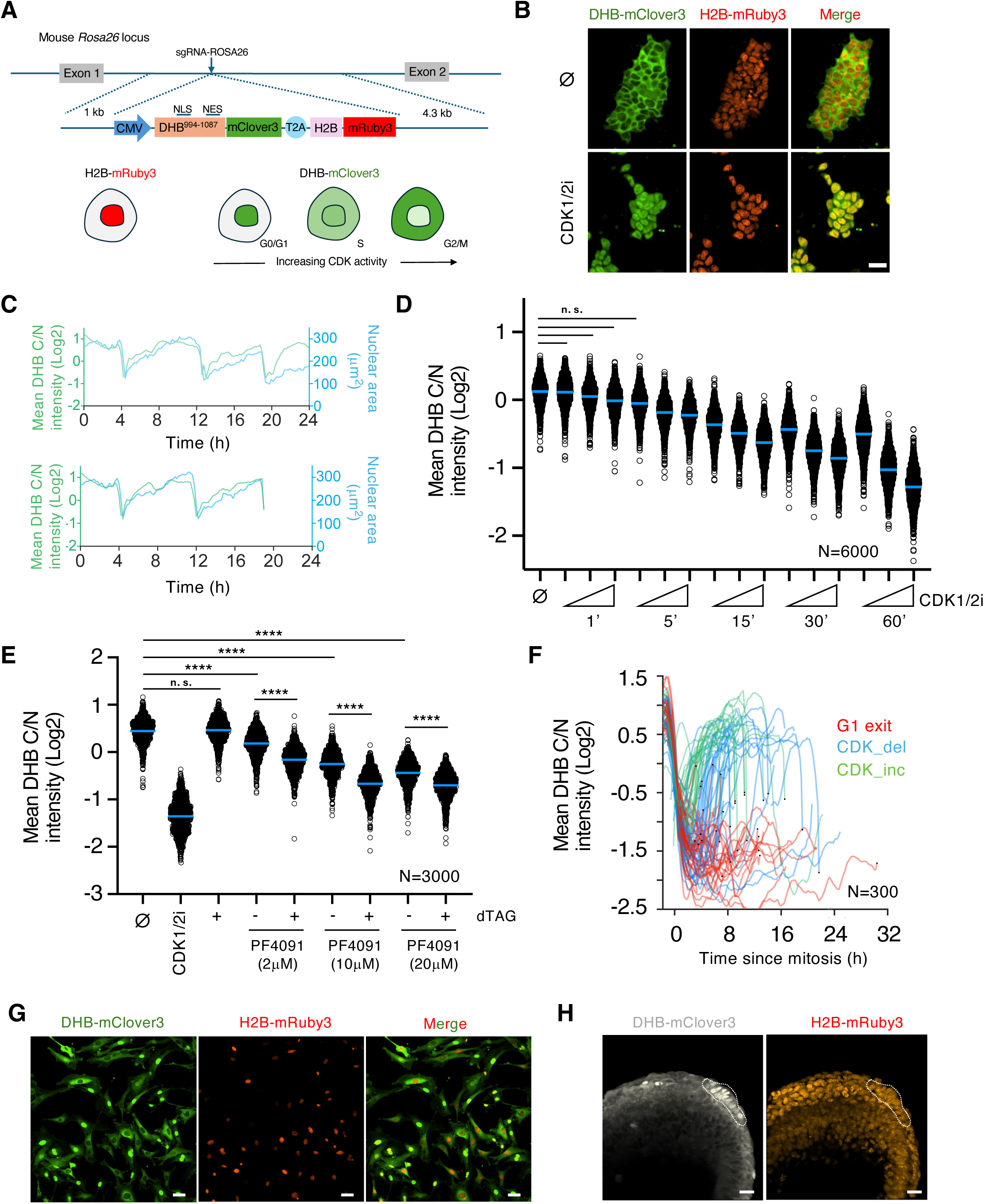
A CDK-KTR transgenic mouse model for live visualization of CDK activity *in vivo*. (A) Schematic representation of the CDK activity sensor targeted at the *Rosa26* locus in ESC. The DNA helicase B (DHB, amino acids 994-1087)-mClover3 KTR shuffles between nucleus and cytoplasm and is used as a readout for CDK activity. NLS: nuclear localization signal. NES: nuclear export signal. (B) Confocal images of mouse ESC^DHB/H2B^ cultures that were untreated (⌀) or CDK1/2i-treated (30μM) for one hour. Scale bar, 30μm. (C) Single-cell CDK activity (based on Cytoplasm/Nuclear (C/N) ratio, green) and nuclear size (blue) traces of representative examples of proliferating ESC^DHB/H2B^. We tracked the same cell (starting point in both plots) and followed daughter and granddaughter cells derived after mitosis. The sudden drop in CDK activity and nuclear size corresponds to a mitotic event. (D) High-throughput imaging (HTI) quantification of C/N mean intensity in untreated (⌀) or CDK1/2i-treated ESC^DHB/H2B^ cultures at different time points with increasing concentration of the inhibitor (1μM, 5μM and 30μM). Center lines indicate mean values. N=6000 cells; All comparisons between untreated and treated cells showed a p-value of p<0.0001 from two-tailed unpaired *t*- tests except those shown as n.s. (non-significant). (E) HTI quantification of C/N mean intensity in untreated (⌀) or dTAG-treated CDK1-degron ESC^DHB/H2B^ cultures combined with the CDK2 inhibitor PF4091 at different concentrations for two hours. Center lines indicate mean values. N=3000 cells; p-values are shown from two-tailed unpaired *t*-tests. **** p<0.0001; n.s.: non-significant. (F) Single-cell CDK activity traces from mouse embryonic fibroblasts (MEF) obtained from ROSA26^DHB/H2B^ mice. All tracks were synchronized in silico to mitosis and the fate of each cell was followed. Criteria to define cells as CDK_inc, CDK_del and G1 exit are specified in Methods. (G) Confocal images of untreated MEF cultures derived from E13.5 ROSA26^DHB/H2B^ mice. Scale bar, 50μm. (H) Confocal images of E7.5 ROSA26^DHB/H2B^ mice showing the location of the node (dashed lines). Scale bar, 20μm.

The DHB-based sensor has been shown to respond primarily to changes in CDK2 activity (Spencer et al, 2013). However, ESC are characterized by elevated and constitutive CDK2 as well as CDK1 activity throughout the cell cycle. Therefore, to determine the contribution of different CDK, we first examined translocation dynamics of the CDK sensor in ESC by using the CDK2 inhibitor PF-07104091 (hereafter PF4091) and observed an inhibitory dose-dependent effect. However, the level of translocation is lower when compared to CDK1/2i-treated ESC^DHB/H2B^ (Figure S1D). Furthermore, CDK2-knockout ESC^DHB/H2B^ with constitutive loss of CDK2 activity showed cytoplasmic localization of the sensor in most of the cells (Figure S1E). This suggests that other CDK may also contribute to the phosphorylation of the reporter. Consistent with previous observations on the minor role of CDK4 in ESC (Savatier et al, 1996; Liu et al, 2017), the CDK4/6 inhibitor PD-0332991(Palbociclib) in combination with PF4091 did not show any effect in promoting nuclear accumulation of the sensor (Figure S1F). To examine the role of CDK1 in regulating the translocation of the CDK reporter, we generated ESC^DHB/H2B^ lines harboring a CDK1 degron (Figure S1G and H). We observed that, although acute CDK1 depletion did not trigger clear nuclear accumulation of the reporter, it showed a dose-dependent cumulative effect on nuclear translocation when combined with PF4091 (Figure 1E). Similar results were observed by using the CDK1 inhibitor RO-3306 in CDK2-knockout ESC^DHB/H2B^ (Figure S1I). These observations demonstrated that translocation dynamics of the CDK sensor are mainly influenced by CDK2 and CDK1 in ESC.

We next microinjected ESC^DHB/H2B^ into blastocysts to generate chimeras and established two independent mouse lines containing the CDK sensor. To test the validity of the mouse model, we isolated mouse embryonic fibroblasts (MEF) and examined the translocation dynamics of the sensor. MEF^DHB/H2B^ showed CDK sensor localization in the nucleus of newly born cells that progressively translocates to the cytoplasm during S-phase (Figure 1F and G). Tracking individual MEF^DHB/H2B^ over time showed the existence of cells undergoing G1 exit as well as two populations of cells defined by their CDK activity kinetics, similar to what has been reported for human fibroblasts (Spencer et al, 2013; Figure 1F and G). Furthermore, we also examined the CDK sensor in cells from the node, a cup-shaped structure that lies on the ventral surface of gastrulating embryos at E7.5 and known to be quiescent. Consequently, we observed that these cells showed nuclear localization of the CDK sensor (Figure 1H). Collectively, we successfully generated a mouse model that can be used to analyze CDK activity during early development.

### CDK activity during pre-implantation development

To examine the CDK activity landscape during pre-implantation development we collected zygotes, 2C, 4C, morula and early, mid, and late-blastocyst stages. Although the signal from the H2B-mRuby3 was visualized as early as in 2C embryos, the signal from the CDK sensor was only detected and reliably quantified from the morula stage onwards (Figure 2A, B and S2A, B). To distinguish between cells contributing to the ICM (embryonic) or the TE (extra-embryonic), we used SOX2 and CDX2 as markers, respectively (Figure 2A-C and S2A-C). Embryonic cells from morula to the mid-blastocyst stage, and independent of their cell lineage, mainly showed cytoplasmic localization of the CDK sensor (Figure 2A-C and S2C). This suggests that elevated CDK activity is driving the first embryonic divisions. However, we consistently observed nuclear accumulation of the CDK reporter in CDX2+ cells at the late blastocyst stage (Figure 2A-C and S2C). To confirm this observation, we performed time-lapse imaging in E3.5 blastocysts cultured *ex-vivo* for additional 20 hours (Figure 2D, Supplementary Video 1). At this stage, most of the cells in the blastocyst showed cytoplasmic localization of the reporter (Figure 2D). However, cells from the TE progressively transitioned to show nuclear localization, especially those from the mural TE, as the embryo approaches implantation (Figure 2D). We occasionally observed TE cells undergoing endoreplication validating previous observations suggesting that this process can occur prior to implantation (Figure S2D).

**Figure 2:**
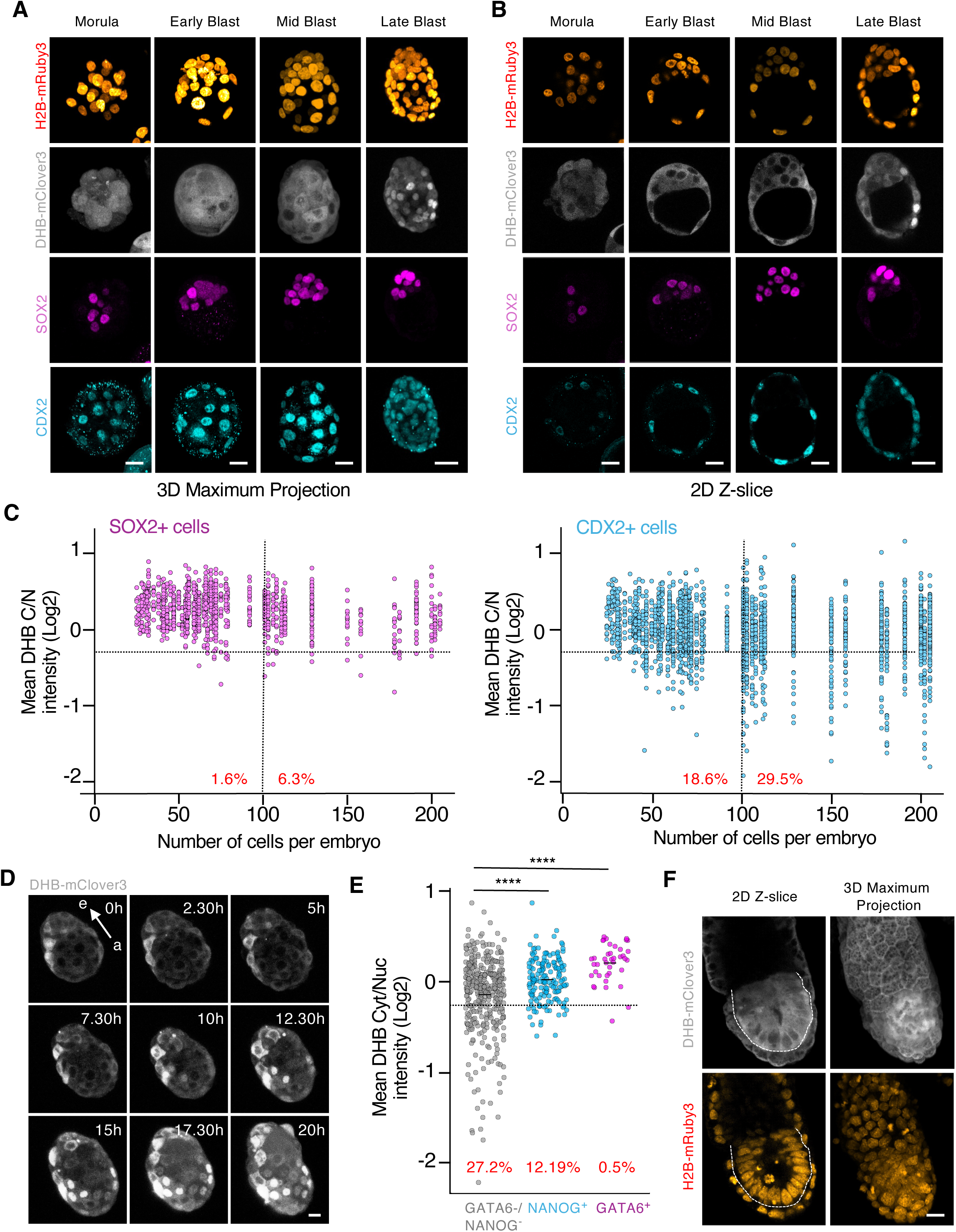
The cells from the TE show a reduction in CDK activity upon implantation. (A and B) Confocal images from hemizygous ROSA26^DHB/H2B^ embryos during pre-implantation development (from morula to late blastocyst). Scale bar, 20μm. (C) Plot showing a quantification of C/N mean intensity in individual cells (each represented by a dot) obtained from hemizygous ROSA26^DHB/H2B^ embryos during pre-implantation development. The collection of dots from the same column belongs to embryos with the same cell count. Embryos were staged based on the number of cells and cells classified as SOX2 (left panel) or CDX2 (right panel) expressing cells in each embryo. In red is the percentage of cells below the defined arbitrary threshold (−0.25) shown for embryos containing 0-100 cells (left) and embryos containing above 100 cells (right). N=54 embryos. (D) Time-lapse microscopy experiment performed in E3.5 isolated hemizygous ROSA26^DHB/H2B^ embryos. One representative embryo is shown. Note how the CDK sensor translocates to the nuclei in TE cells. Arrow indicates the embryonic (e)- abembryonic (a) axis. Scale bar, 15μm. (E) Plot showing a quantification of C/N mean intensity in individual cells obtained from a pool of hemizygous E4.5 ROSA26^DHB/H2B^ embryos. In red is the percentage of cells below the arbitrary defined threshold (−0.25). p-values are shown from two-tailed unpaired *t*-tests. **** p<0.0001; N=5 embryos. (F) Confocal images from E6.5 hemizygous ROSA26^DHB/H2B^ embryos. Dashed line surrounds the EPI. Scale bar, 20μm.

We also examined the levels of CDK activity in GATA6+ cells (PrE) and NANOG+ cells (EPI) by analyzing E4.5 pre-implantation blastocysts and observed cytoplasmic localization of the CDK sensor (Figure 2E and S2E). Fast proliferation kinetics in the PrE are needed after implantation to quickly expand and generate the visceral and parietal endoderm (VE and PE, respectively). This contrasts with the nuclear localization of the CDK reporter observed in NANOG-/GATA6- cells which correspond to TE cells (Figure 2E).

Finally, we examined the level of CDK activity in cells from the post-implantation epiblast by isolating E5.5 (Figure 2F). These cells majorly showed cytoplasmic localization of the CDK sensor in agreement with the fast proliferation kinetics of the post-implantation epiblast (Figure 2F). In summary, our data revealed that embryonic cells maintain elevated levels of CDK activity to sustain the high proliferation rate needed while cells from the TE downregulate CDK activity prior to implantation.

### FGF4-dependent regulation of CDK activity in the TE

To understand the mechanisms underlying the differential regulation of CDK activity between embryonic and extra-embryonic cells, we used ESC and trophoblast stem cells (TSC), as the corresponding *in vitro* counterparts of the ICM and TE, respectively. TSC^DHB/H2B^ showed cytoplasmic localization of the CDK sensor, rapid cell cycle dynamics and respond to CDK inhibition similarly to ESC (Figure 3A and S3A and B). This suggests that CDK activity downregulation in TE cells depends on non-autonomous mechanisms. Thus, we next examined the response of the sensor to the deprivation of self-renewal signals. ESC self-renew in the presence of leukemia inhibitory factor (LIF) (Williams et al, 1988) while TSC rely on fibroblast growth factor 4 (FGF4) to sustain their functional properties (Tanaka et al, 1998). Indeed, FGF4 deprivation induces differentiation into non-proliferating TGC following p57^KIP2^-dependent CDK1 inhibition (Ullah et al, 2008). LIF withdrawal in ESC, which promotes differentiation and exit from naïve pluripotency (Kalkan et al, 2017), did not induce nuclear translocation of the DHB reporter or changes in proliferations dynamics (Figure 3A). However, FGF4 withdrawal resulted in most TSC showing nuclear localization of the CDK sensor (Figure 3A). We tracked single ESC and TSC to examine their individual behavior. In the presence of LIF, ESC showed a homogenous behavior with cells robustly cycling, which was maintained upon LIF withdrawal (Figure 3B). However, we identified cells with different CDK activity dynamics and cells exiting in G1 phase in cultures of FGF4-treated TSC (Figure 3B). Moreover, in FGF4-deprived cultures of TSC we identified cells exiting from G1 or G2 phases as well as cells undergoing endoreplication (Figure 3B and S3C). Consistently, we observed that, whereas differentiated TSC with nuclear CDK sensor showed elevated levels of the CDK inhibitor p57^KIP2^, ESC did not show changes in p57^KIP2^ levels (Figure S3D and E). In summary, TSC were intrinsically more prone than ESC to downregulate CDK activity upon changes in the levels of self-renewing signals.

**Figure 3:**
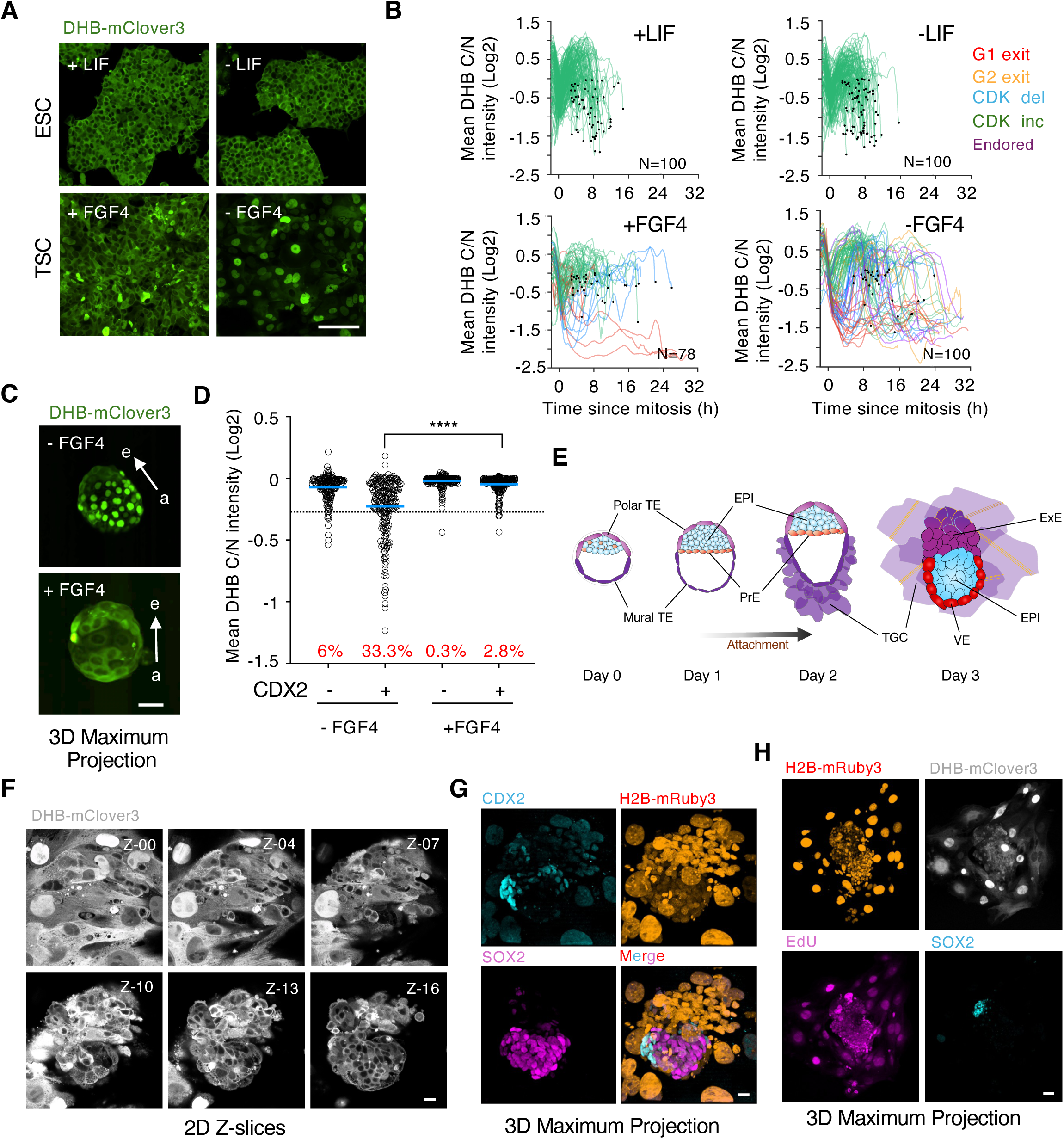
ICM-derived FGF4 establishes a signaling gradient through the embryonic- abembryonic axis. (A) Confocal images of TSC^DHB/H2B^ and ESC^DHB/H2B^ cells cultured with or without self-renewal signals for seventy-two hours. Scale bar, 100μm. (B) Single-cell CDK activity traces from TSC^DHB/H2B^ and ESC^DHB/H2B^ cells cultured with or without self-renewal signals for seventy-two hours. All tracks were synchronized in silico to mitosis and the fate of each cell was followed. Criteria to define cells as CDK_inc, CDK_del, Endored, G1 exit and G2 exit are specified in Methods. (C) Confocal images of representative untreated or FGF4-treated hemizygous ROSA26^DHB/H2B^ embryos cultured for 20 hours after isolation at E3.5. Scale bar, 50μm. Arrows indicate the embryonic (e)- abembryonic (a) axis. (D) Plot showing quantification of C/N mean intensity in a pool of CDX2+ or CDX2- cells from untreated (N=5 embryos) or FGF4-treated (N=9 embryos). P-value is shown from a two-tailed unpaired *t*-test. **** p<0.0001. In red is the percentage of cells below the defined arbitrary threshold (−0.25). (E) Schematic representation of the embryonic development *in vitro* of a E3.5 (Day 0 for *in vitro* culture) blastocyst beyond implantation stages. TE: trophectoderm; EPI: epiblast; PrE: primitive endoderm; TGC: trophoblast giant cell; VE: visceral endoderm; ExE: extraembryonic endoderm. (F) Series of 2D-Z confocal images (3μm slice interval) of a representative hemizygous ROSA26^DHB/H2B^ embryo cultured *in vitro* three days after isolation at E3.5. Scale bar, 20μm. (G) Confocal images of the same hemizygous ROSA26^DHB/H2B^ embryo from (F). Scale bar, 20μm. (H) Confocal images of a representative hemizygous ROSA26^DHB/H2B^ embryo cultured three days after isolation at E3.5. Scale bar, 50μm.

We investigated whether the lack of FGF4 signaling in TE cells *in vivo* was responsible for the drop in CDK activity prior to implantation. Since FGF4 is exclusively expressed by the cells from the ICM (Boroviak et al, 2015), the levels of FGF4 might not be enough to sustain elevated levels of CDK activity in cells from the TE, especially those distant from the ICM. Thus, we isolated E3.5 blastocysts and cultured them for additional 20 hours with or without exogenous FGF4. Addition of FGF4 prevented the accumulation of the CDK sensor in the nuclei of TE cells demonstrating the existence of an FGF4-dependent signaling gradient through the embryonic-abembryonic axis (Figure 3C and D, Supplementary Video 2).

We next examined *in vivo* the differentiation process of the embryo during peri-implantation and onwards. Following implantation, the polar TE expands and generates the ExE, trophoblast progenitor cells that will later differentiate to form the embryonic part of the placenta. In addition, the PrE gives rise to the PE which migrates over the mural TE, and to the VE which surrounds the ExE and the EPI. Finally, the EPI organizes into a rosette of polarized cells and undergo a process of lumenogenesis to form the egg cylinder. To visualize these steps, we used a protocol that supports the *in vitro* development of mouse blastocysts beyond implantation stages (Bedzhov et al, 2014; Figure 3E). This protocol requires the sequential use of two different media, IVC1 which triggers TE differentiation into TGC and promotes adhesion to the plate within the first 24 hours, and IVC2 which supports the growth of the emerging egg cylinder. We isolated E3.5-E3.75 blastocysts and imaged them. Following attachment, cells from the mural TE quickly spread and differentiated into TGC (Figure 3F and G, Supplementary Video 3). This process could be visualized by the progressive increase in nuclear size following endoreplication. Indeed, differentiated TGC showed consecutive rounds of nuclear/cytoplasmic shuttling of the CDK sensor and EdU incorporation with no signs of mitotic events (Figure 3H and S4A). To assess whether genome duplication in cells undergoing endoreplication was driven by CDK2, we added CDK2i and observed that CDK2i-treated embryos did not show the expected rounds of nuclear/cytoplasmic shuttling of the CDK sensor (Figure S4B). This observation demonstrated that, while CDK1 needs to be inhibited to enable endoreplication, CDK2 drives genome replication in TGC. Furthermore, the attached embryos also expressed p57^KIP2^ which, like FGF4-deprived TSC, might keep CDK1 inhibited to enable endoreplication (Figure S4C). Interestingly, we did not detect similar expression of p57^KIP2^ in TE cells from pre-implantation embryos suggesting that implantation is required to trigger p57^KIP2^ expression *in vivo* (Figure S4D).

Embryonic cells identified by SOX2 expression were characterized by cytoplasmic localization of the CDK reporter, consequent with their high proliferation rate (Figure 3F and G). Furthermore, cells from the polar TE, which will further expand to generate more TGC, were found adjacent to embryonic cells and showed cytoplasmic localization of the sensor (Figure 3F and G). We also cultured embryos into the stage of egg cylinder and observed that embryonic cells from the epiblast, the ExE and the VE still contained high levels of CDK activity to sustain the elevated rate of cell proliferation needed prior to gastrulation (Figure S4E). Our findings revealed an FGF4- dependent embryonic-abembryonic axis that enabled TE cells to downregulate CDK activity and prime them toward TGC differentiation.

### Human TE-like cells show nuclear accumulation of the CDK sensor in a gastruloid model

We next investigated the level of conservation in regulating CDK activity in TE cells by examining human cells. Thus, we established human ESC (hESC) lines constitutively expressing the CDK sensor (hESC^DHB/H2B^). Although hESC are in a primed pluripotent state, a more advanced developmental stage compared to naïve mouse ESC, they also showed cytoplasmic localization of the CDK sensor and confirmed the elevated levels of CDK activity (Figure 4A and S5A). Moreover, naïve hESC^DHB/H2B^ lines also showed similar levels of CDK activity (Figure S5B). hESC^DHB/H2B^ also responded to the CDK1/2i as well as to the acute inhibition of CDK2 using PF4091 (Figure S5A and B). Furthermore, by examining CDK2-deficient hESC^DHB/H2B^ we confirmed that the DHB/H2B reporter was also regulated by CDK1 in the absence of CDK2 (Figure S5C) (Aleem et al, 2005). Individual tracking of hESC^DHB/H2B^ revealed, that following mitosis, CDK activity was maintained over the cell cycle (Figure 4B). To examine the fluctuations of CDK activity during TE specification, we used a 2D system to mimic human gastrulation (Warmflash et al, 2014; Minn et al, 2020). In this system, circular-shaped micropatterned hESC, stimulated with BMP4, differentiate into radially organized cellular rings containing the three germ layers. Starting from the outer ring of cells toward the center of the circle, cells were specified as trophectoderm, endoderm, mesoderm and ectoderm clearly defined by specific markers for each germ layer (Figure S5D-G). Therefore, it is an ideal system to analyze the localization of the CDK sensor upon cell fate specification. Interestingly, we observed an enrichment in cells showing nuclear accumulation of the reporter in the outer ring of TE-like cells, characterized by containing the population of CDX2+ cells and a high level of BMP signaling (Figure 4C and S5D-G). This was specific to BMP4-treated gastruloids as untreated cells neither properly differentiated nor showed nuclear accumulation of the DHB reporter across the gastruloid (Figure 4C). To quantify the percentage and location of cells showing nuclear signal of the CDK sensor, we first divided the gastruloid into 5 circular bins of approximately 100μm and plotted the DHB ratio for a defined number of total cells within the gastruloid (Figure 4D and S6A). We detected a clear accumulation of cells with nuclear signal in the outer bin of BMP4-treated gastruloids suggesting that the population of TE-like cells was enriched in cells with low levels of CDK activity (Figure 4D). Thus, we next examined the level of DHB nuclear accumulation present in TE-like cells, defined by the expression of CDX2, and compared those to CDX2- cells. We detected a clear increase in the number of cells with nuclear accumulation of the CDK sensor in CDX2+ cells (Figure 4E). As expected, these cells were located at the outermost part of the gastruloid (Figure 4F and S6B). We also confirmed this increase in TE-like cells by examining the levels of translocation in TFAP2C-positive cells, a different marker labeling TE-like cells (Figure S6C and D). Importantly, cells expressing markers from the mesoderm or the ectoderm (Brachyury or SOX2, respectively), did not contain cells with detectable sensor in the nucleus (Figure S6E-G). Finally, we examined whether human TE-like cells induced the expression of p57^KIP2^ to limit CDK activity in these cells. Indeed, we detected enrichment of p57^KIP2^-expressing cells in the outer ring of differentiated gastruloids (Figure S6H). In summary, these results showed how TE-like cells down-regulate CDK activity in gastruloids underscoring the existence of conserved regulation in this process in humans.

**Figure 4:**
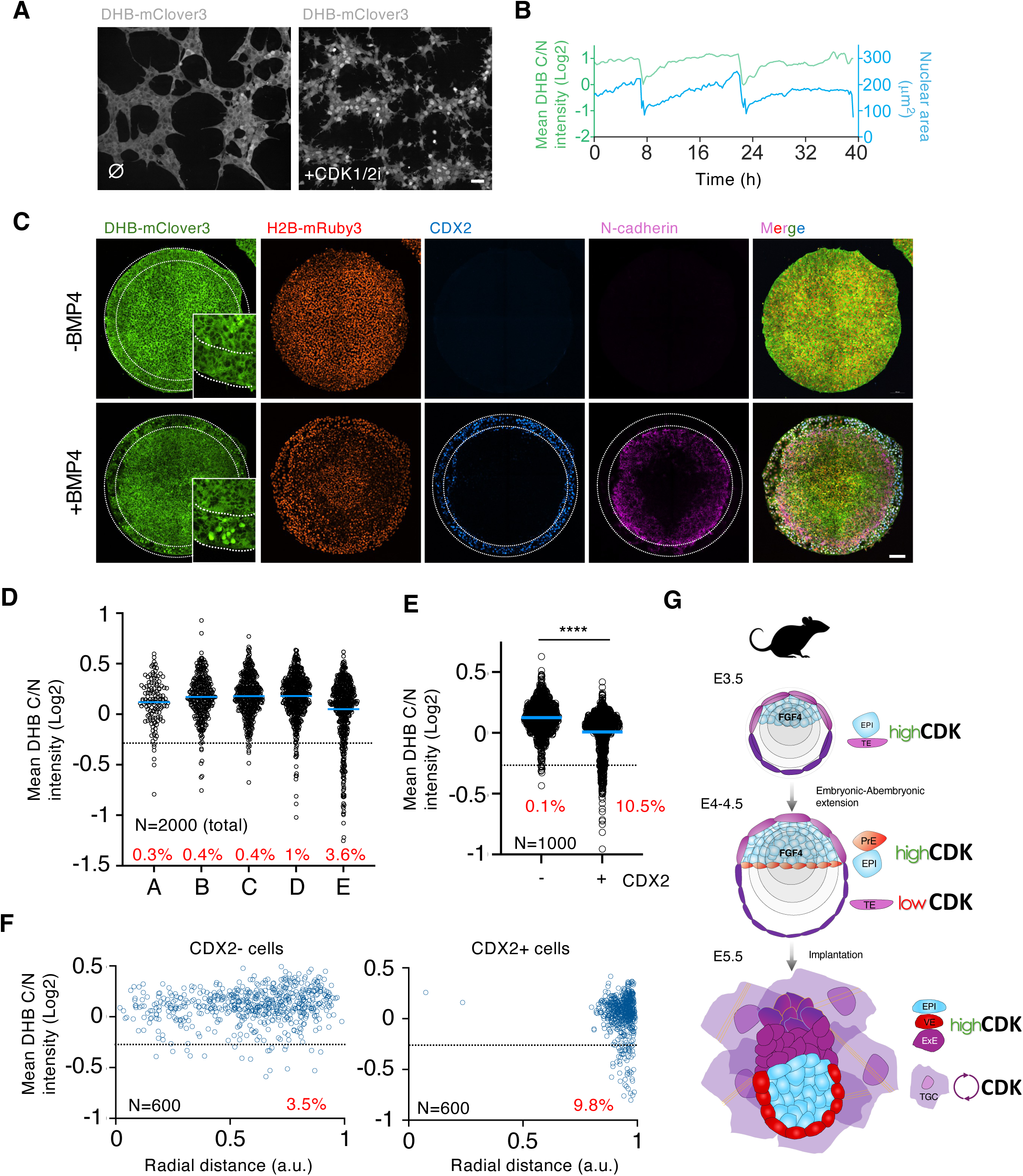
Human TE-like cells show nuclear translocation of the CDK sensor. (A) Confocal images of human ESC^DHB/H2B^ cultures that were untreated (⌀) or CDK1/2i-treated (30μM) for one hour. Scale bar, 50μm. (B) Single-cell CDK activity and nuclear trace of a representative proliferating human ESC^DHB/H2B^. The sudden drop in CDK activity and nuclear size corresponds to a mitotic event. (C) Confocal images of representative gastruloids uninduced or BMP4-induced for seventy-two hours. Insets can be found for DHB-mClover3 with higher magnification. Dashed lines indicate the outer ring of TE-like cells identified by CDX2 expression. Scale bar, 100μm. (D) Plot showing a quantification of C/N mean intensity in a total number of 2000 random individual cells distributed according to their position in defined rings from the center of the gastruloid (see Figure S8A for details). In red is the percentage of cells per ring below the defined arbitrary threshold (<-0.25). Data was obtained by combining multiple gastruloids from the same experiment (N=12 gastruloids). (E) Plot showing a quantification of C/N mean intensity in 1000 random CDX2 positive or negative cells from human gastruloids. p value is shown from a two-tailed unpaired *t*-test. **** p<0.0001. In red is the percentage of cells below the defined arbitrary threshold (−0.25). Data was obtained by combining multiple gastruloids from the same experiment (N=12 gastruloids). (F) Plot showing a quantification of C/N mean intensity in 600 random CDX2+ or CDX2- cells distributed according to their radial distance from the center of the gastruloid (see Figure S8A for details). In red is the percentage of cells per ring below the defined arbitrary threshold (<-0.25). Data was obtained by combining multiple gastruloids from the same experiment (N=12 gastruloids). (G) Schematic model summarizing our findings. TE: trophectoderm; EPI: epiblast; PrE: primitive endoderm; TGC: trophoblast giant cell; VE: visceral endoderm; ExE: extraembryonic endoderm.

## DISCUSSION

Early embryonic development is characterized by fast proliferation dynamics. This high proliferation rate is recapitulated in ESC and depends on constitutive and elevated levels of CDK activity throughout the cell cycle associated with low or non-detectable levels of CDK inhibitors. These observations suggested that cells from the ICM may regulate CDK activity in a similar manner. However, whether there are lineage-specific changes in CDK activity during early development or whether they can influence cell fate *in vivo* remained unclear. By using a mouse model containing a CDK-KTR live-cell sensor, our results revealed the existence of a dynamic and lineage-specific CDK activity landscape during early embryonic development (Figure 4G).

We showed how cells from the TE progressively lower CDK activity as the mouse blastocyst approaches implantation. This switch is induced by decreased FGF4 availability and not by TE-specific autonomous mechanisms as the addition of exogenous FGF4 prevented the drop in CDK activity. Our results suggested that FGF ligands preferentially act locally and establish a signaling gradient through the embryonic-abembryonic axis. Similar observations were made by using an ERK-KTR mouse model (Simon et al, 2020; Pokrass et al, 2020). FGF4 signaling is required for maintaining TE identity and proliferation in TSC suggesting that low FGF4 signaling in the mural TE primes these cells to differentiate. Conversely, FGF4 signaling is required for cells of the polar TE to retain multipotency and high levels of CDX2 upon implantation (Christodoulou et al, 2019). In summary, FGF ligands produced by the ICM act as positional cues to spatially and temporally pattern CDK activity in the pre-implantation embryo.

Using an *in vitro* protocol that supported embryo development beyond implantation stages, we visualized the differentiation process of mural TE cells to TGC. Mural TE cells in the blastocyst showed nuclear accumulation of the CDK sensor indicating they were either arrested in G1 or in G2 prior to attachment. Embryo attachment induced the entry of mural TE cells into endocycles suggesting that mechanical signals are the trigger of this process. Similar to differentiating TSC *in vitro*, TGC also upregulated the expression of p57^KIP2^ *in vivo*. This upregulation leads to CDK1 inhibition, which prevents mitosis, while the levels of CDK2 activity oscillate to enable genome replication. Although we could visualize endoreplication in peri-implantation blastocysts, these events were rare. In agreement, we did not detect significant p57^KIP2^ expression prior to implantation.

High levels of CDK activity in ESC are associated with their pluripotency potential and self-renewal abilities (Wang et al, 2017; Michowski et al, 2020). Indeed, a lengthening in the duration of the cell cycle through modulation of CDK activity leads to differentiation (Pauklin and Vallier, 2013). Furthermore, differentiation in ESC is accompanied by changes in cell cycle structure by imposing CDK cyclic activity (White et al, 2005). Moreover, overexpression of specific CDK or cyclins in somatic cells facilitate reprogramming to induced pluripotent stem cells (Ruiz et al, 2011). These observations highlight the intimate link between CDK activity and cell fate in pluripotent cells. We began to detect changes in CDK activity in TE cells from E3.5 blastocysts. However, at this stage the blastocyst is already segregated into two distinct cell lineages, TE and ICM. Similarly, subsequent segregation of the ICM into EPI and PrE is characterized by the absence of clear changes in CDK activity as most EPI and PrE cells show cytoplasmic localization of the CDK sensor. Thus, our results indicate that cell fate decisions during pre-implantation development are not determined by changes in CDK activity. Nonetheless, we cannot discard that manipulating levels of CDK activity and/or altering cell cycle dynamics could influence the efficiency or the balance of specific cell fate decisions.

Our results also revealed lineage-specific regulation of CDK activity in human cells pointing to conserved regulatory mechanisms in mammals during implantation. Indeed, human TE-like cells in an *in vitro* system of gastrulation showed nuclear accumulation of the CDK sensor compared to cells from other germ layers. Although murine and primate embryos are morphologically similar around implantation, they begin to show different behavior and morphogenic transformations at implantation (Shahbazi and Zernicka-Goetz, 2018). The mouse blastocyst implants by the mural TE and differentiates into TGC while the polar TE generates the ExE. On the contrary, the human embryo uses the TE cells from the embryonic pole to implant and transforms into villous cytotrophoblasts, a bipotent TE stem cell progenitor. The different behavior of the TE cells during peri-implantation seems to be, at least partially, responsible for the different shapes of the post-implantation embryo (Christodoulou et al, 2019). Nevertheless, besides the different fate of TE cells after implantation in mouse and human embryos, TE-like cells differentiated in gastruloids show a transcriptional signature that corresponds to TE cells from the E7 pre-implantation human blastocyst (Minn et al, 2020) and thus, seem comparable to TE cells from mouse pre-implantation embryos.

Finally, based on the excellent dynamic range of the CDK sensor to visualize changes during pre-implantation development, we envision that this reporter mouse model will be broadly applicable to studying CDK signaling dynamics in other physiological or non-physiological contexts including cancer.

## ACKNOWLEDGMENTS

We thank David Santamaria, Carmen J. Williams and members from the Laboratory of Genome Integrity, for the helpful comments and discussion on this work. We also thank Ferenc Livak, Shafi Siddiqui and the CCR Flow cytometry Core, Gianluca Pegoraro and the High-Throughput Imaging Facility, the Microscopy Core and the HPC Biowulf cluster for experimental support at the NCI. Research in S.R. laboratory is supported by the Intramural Research Program of the NIH.

## AUTHOR CONTRIBUTIONS

B.S. and S.R. conceived the study. B.S. and S.R., designed, performed, and analyzed most of *in vitro* and *in vivo* experiments. J.A.C. and S.D.C. performed live imaging and tracking experiments *in vitro*. A.D.T. and M.J.K. analyzed confocal microscopy data. M.A.C. provided technical support. P.R.S. and N.Y.M. provided help with PDMS stamps for gastruloid experiments. S.R. supervised the study and wrote the manuscript with comments and help from all authors.

## METHODS

| REAGENT | SOURCE | IDENTIFIER |
| --- | --- | --- |
| **Antibodies** |  |  |
| Mouse Anti-CDX2 | BioGenex | MU392A-UC, RRID:AB_2923402 |
| Rabbit Anti CDX2 | Abcam | ab76541, RRID:AB_1523334 |
| Mouse Anti p57<sup>KIP2</sup> | Santa Cruz | sc-56341, RRID:AB_785045 |
| Mouse Anti TFAP2C | Santa Cruz | sc-12762, RRID:AB_667770 |
| Rabbit Anti Brachyury | Abcam | ab209665, RRID:AB_2750925 |
| Rabbit Anti Sox2 | Cell Signaling | 23064S, RRID:AB_2714146 |
| Rabbit Anti Phospho-CDK substrate Motif | Cell Signaling | 9477, RRID:AB_2714143 |
| Goat Anti Gata6 | R&D systems | AF1700, RRID:AB_2108901 |
| Rabbit Anti Nanog | Abcam | ab80892, RRID:AB_2150114 |
| Mouse Anti N-Cadherin | Cell Signaling | #14215, RRID:AB_2798427 |
| Rabbit Anti Phospho-SMAD1/5 | Cell Signaling | #9516, RRID:AB_491015 |
| Rabbit Anti CDK1 | Millipore | 06-923, RRID:AB_11214395 |
| **Chemicals, peptides and recombinant proteins** |  |  |
| CDK1/2 inhibitor | Sigma-Aldrich | 217714 |
| CDK2 inhibitor (PF-07104091) | Axon MEDCHEM | 3753 |
| CDK1 inhibitor (Ro3306) | Sigma-Aldrich | SML0569 |
| Palbociclib (PD-0332991) | Selleckchem | S1116 |
| Recombinant Human FGF4 | R&D Systems | 235-F4-025 |
| Heparin | Sigma-Aldrich | H3149-10KU |
| M2 | EMD Millipore | MR-015-D |
| EmbryoMax KSOM | EMD Millipore | MR-106-D |
| Silicon oil | Chem-Cruz | sc-215854 |
| GoTaq® Hot Start Polymerase | Promega | M500B |
| Proteinase K | Meridian Bioscience | BIO-37084 |
| Paraformaldehyde (PFA) | EMS | 15710 |
| Acid Tyrode's | Sigma-Aldrich | T1788 |
| Glycine | Sigma-Aldrich | 50046-250G |
| Triton X-100 | Sigma-Aldrich | T8787 |
| DMEM | Gibco | 11965-092 |
| DMEM-F12 | Gibco | 11320-033 |
| RPMI | Quality Biological | 112-025-101 |
| NEAA | Gibco | 11140-050 |
| Glutamax | Gibco | 35050-61 |
| Sodium Pyruvate | Gibco | 11360-070 |
| 2-mercaptoethanol | Gibco | 31350-010 |
| Fetal Bovine Serum | GEMINI Bio products | 100-525 |
| 0.05% Trypsin EDTA | Gibco | 25300-054 |
| Gelatin | Sigma-Aldrich | G1890 |
| Penicillin/Streptomycin | Gibco | 15070-063 |
| mTeSR PLUS | Stem Cell Technologies | 100-0276 |
| RSeT™ Medium | Stem Cell Technologies | 05978 |
| Dispase | Stem Cell Technologies | 7923 |
| Accutase | Gibco | A11105-01 |
| Matrigel Growth Factor Reduced | Corning | 3566231 |
| Rho kinase inhibitor, Y27632 | Selleckchem | S1049 |
| BMP4 | R&D Systems | 314-BP-010 |
| Laminin | BioLamina | LN521-04 |
| ITS-X | Life Technologies | 51500-056 |
| β-Estradiol | Sigma-Aldrich | E8875 |
| Progesterone | Sigma-Aldrich | P0130 |
| N-Acetyl-L-cysteine | Sigma-Aldrich | A7250 |
| Pregnant Mare Serum Gonadotropin | PROSPEC | hor-272 |
| Chorionic Gonadotropin Human | PROSPEC | hor-250 |
| Bovine Serum Albumin (BSA) | Sigma-Aldrich | 3117332001 |
| Tween 20 | Sigma-Aldrich | 9005-64-5 |

**Supplementary Table 1.**
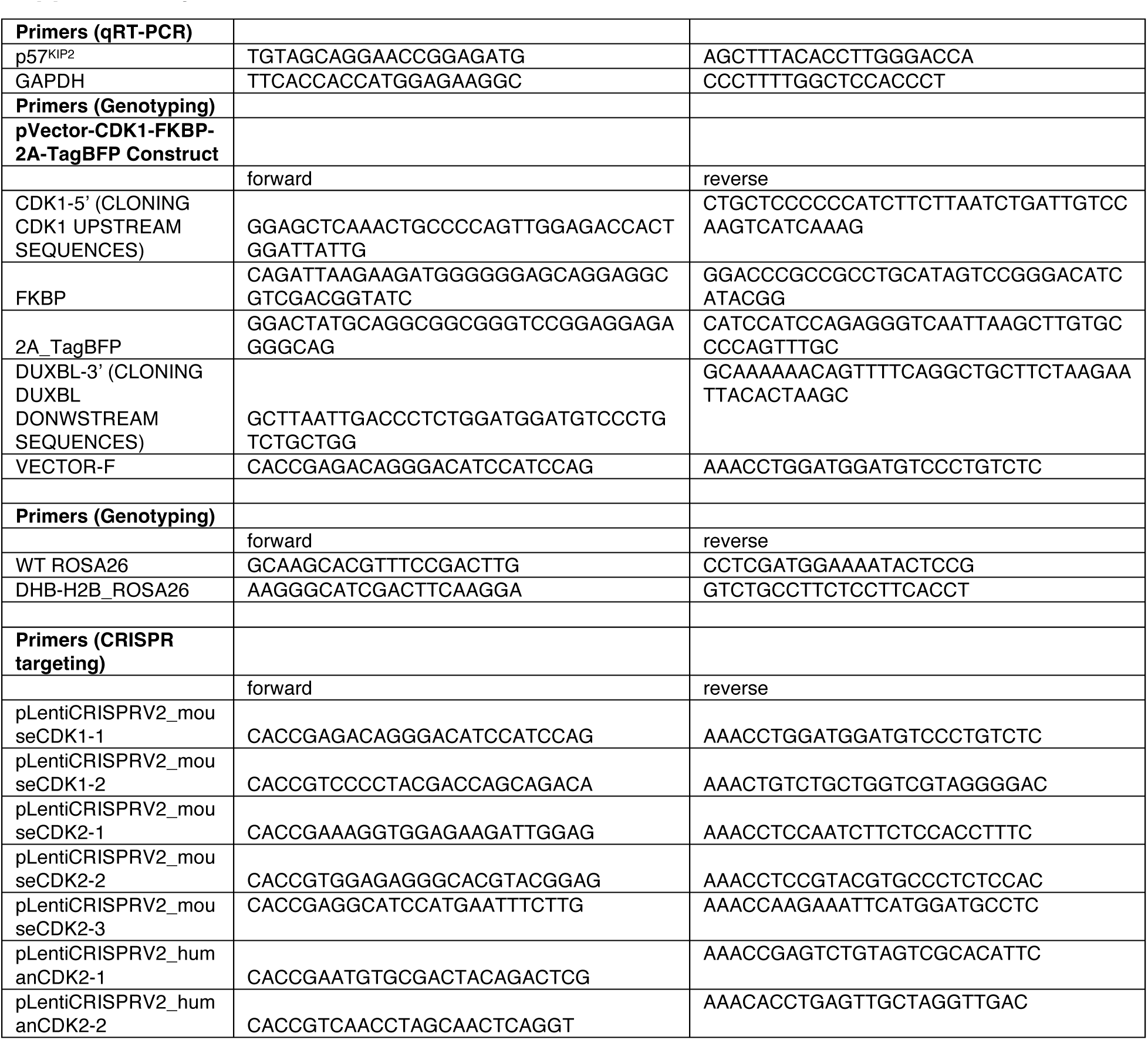

### Cell lines and culture

Wild-type (R1) ESC were grown on 0.1% gelatin-coated plates or alternatively on a feeder layer of growth-arrested MEF in high-glucose DMEM (11965-092, Gibco) supplemented with 15% FBS, 1:500 leukemia inhibitory factor (LIF; made in house), 0.1 mM nonessential amino acids (11140-050, Gibco), 1% glutamax (35050-61, Gibco), 55 mM β-mercaptoethanol (31350-010, Gibco) and 1% penicillin/streptomycin (15070-063, Gibco) at 37°C and 5% CO2. Cells were routinely passaged with Trypsin 0.05% (25300-054, Gibco). Media were changed every other day and passaged every 2–3 days. MEF were grown in DMEM, 10% FBS, and 1% penicillin/streptomycin. HEK293T (a gift from Dr. Inder Verma, SALK Institute, San Diego, CA, USA) cells were grown in DMEM, 10% FBS, and 1% penicillin/streptomycin. Generation of infective lentiviral particles and ESC infections were performed as described (Ruiz et al, 2011).

To generate ESC^DHB/H2B^ lines, we first made a targeting construct using CRE-treated and BstBI/MluI digested LSL-Cas9-ROSA26TV plasmid donor (gift from Feng Zhang, #61408, Addgene) to subclone the DHB-mClover3_2A_H2B-mRuby3 fragment. This fragment was generated by Gibson assembly (New England Biolabs) using synthetized gene fragments (IDT technologies). To generate CDK1^FKBP/FKBP^ ESC lines, we followed a similar approach. Fragments to generate the recombination construct (2xHA-FKBP-2A-TagBFP) were amplified by PCR and ligated them by Gibson cloning using the plasmid backbone from CTCF-mAC donor (gift from Masato Kanemaki, #140646, Addgene). ESC were co-transfected using jetPRIME (PolyPlus transfection) with the targeting vector as well as with LentiCRISPRv2 (gift from Feng Zhang, #52961, Addgene) encoding a sgRNA designed to target the ROSA26 or CDK1. Targeting events are uncommon and therefore, mClover3+ or TagBFP ESC were sorted on a BD FACSAria Fusion instrument.

Trophoblast stem cells (TSC) (gift from Magdalena Zernicka-Goetz, Caltech) were cultured on an irradiated layer of MEF in RPMI-1640 medium supplemented with 20% FBS, Glutamax (1x, Gibco), 0.1 mM nonessential amino acids (1x, Gibco), Sodium pyruvate (1x, Gibco), 55 mM 2-Mercaptoethanol (100 μM, Gibco), 1% penicillin/streptomycin (Gibco), FGF4 (25 ng/mL), and heparin (1 μg/mL). To generate TSC^DHB/H2B^ lines, we first generated a lentiviral vector to constitutively express the transgene DHB-mClover3_2A_H2B-mRuby3 by cloning this fragment into BamHI/EcoRI digested FUGW (Gift from David Baltimore, #14883, Addgene). TSC were trypsinized and infected in suspension with infective lentiviral particles. Infected clonal TSC lines were established and amplified in culture.

Human ESC (hESC, H9 cell line, Wicell Institute) were cultured in wells coated with growth factor reduced Matrigel (354230, Corning) in mTeSR plus medium (Stemcell, #100-0276) with daily changes of media. Cells were passaged at 80% confluency using 1mg/mL Dispase (Stemcell, #07923) and manual scrapping and cultured at 37°C and 5% CO2. To generate hESC^DHB/H2B^ lines, H9 cells were trypsinized and infected in suspension with infective lentiviral particles. Infected clonal hESC lines were established and amplified in culture. To reprogram primed H9 cells into a naïve state we used RSeT medium (Stem Cell Technologies) following manufacturer’s instructions. All experiments conducted with hESC followed protocols approved by the National Institutes of Health ESCRO committee.

None of the cell lines used in this manuscript are listed in the International Cell Line Authentication Committee (ICLAC) Database of Cross-contaminated or Misidentified Cell Lines. Testing for mycoplasma contamination in our cells was performed routinely.

### Mice

All mouse breeding and experimentation followed protocols approved by the National Institutes of Health Institutional Animal Care & Use Committee (ACUC, protocol 07-37) Guideline for Breeding and Weaning. C57BL/6J mice obtained from the Jackson Laboratory, Bar Harbor, ME. Mice were maintained in a dark/light cycle of 12 hours each in a temperature range of 68°-76°F and a range of 30-70% humidity. To generate the ROSA26^DHB/H2B^ mice, we microinjected ESC^DHB/H2B^ into C57BL/6 blastocysts and transferred to pseudopregnant females. Several ROSA26^DHB/H2B^ mouse chimeras were generated from which we established two mouse lines. The two independent ROSA26DHB^/H2B^ mouse lines generated were functionally comparable although all experiments shown in this manuscript correspond to one of these lines. ROSA26^DHB/H2B^ mice were generated by the Mouse Modeling & Cryopreservation (MMC) facility at the National institutes of health (NIH). To genotype, genomic DNA was extracted from a small ear biopsy and PCR was performed with GoTaq® Hot Start Polymerase (Promega) using specific primers (Supplementary Table 1).

### Mouse embryonic fibroblast isolation

Mouse embryos from pregnant females were collected at E13.5. The head and organs from the embryos were removed in PBS and thereafter transferred into a clean dish. Embryos were finely minced in minimal amount of carry over PBS and incubated for 15 min at 37°C with 1mL of 0.05% trypsin/EDTA. Cells were dissociated by energic pipetting up and down. Trypsin was then inactivated in DMEM containing 10% FBS and 0.1 mM nonessential amino acids (Gibco). Cells from an individual embryo were centrifuged, plated in a p100 plate and routinely passaged every 3-4 days.

### Embryo collection and culture

For embryo collection, male heterozygous ROSA26^DHB/H2B^ mice were mated with four to six weeks old female wild-type mice. Females were intraperitoneally injected with 5 IU of pregnant mare serum gonadotropin (PMSG, Prospec), followed 48 hours later by 5 IU of human chorionic gonadotropin (hCG; Sigma-Aldrich). Alternatively, eight-weeks-old naturally pregnant females were also used for embryo isolation. Pre-implantation embryos (zygotes, 2C, 4C-stage embryos, morulas and blastocysts of different age) were flushed from the oviducts and collected in M2 media (MR-015-D, Sigma-Aldrich) while post-implantation embryos (E5.5 and E7.5) were collected in DMEM/F-12 (Gibco) supplemented with 10% FBS, 15 mM HEPES, 1% Penicillin-Streptomycin and 1X L-glutamine. For time-lapse microscopy experiments performed with isolated E3.5 hemizygous ROSA26^DHB/H2B^ embryos we first digested the zona pellucida with Tyrode’s acid, washed three times in M2 and then transferred to IVC1 medium (DMEM/F12 supplemented with 20% (vol/vol) heat-inactivated FBS, 2 mM L-glutamine, penicillin/streptomycin (25 μg/ml), 1× ITS-X (10 mg per liter insulin, 5.5 mg per liter transferrin, 0.0067 mg per liter sodium selenite and 2 mg per liter ethanolamine), 8 nM β-estradiol, 200 ng/ml progesterone and 25 μM *N*-acetyl-L-cysteine) for up to 48 hours in 18-wells chambered coverslips (Ibidi). Following attachment and spread, embryos were switched and further cultured in IVC2 (DMEM/F12 supplemented with 30% (vol/vol) KSR, 2 mM L-glutamine, penicillin (25 units/ml)/streptomycin (25 μg/ml), 1× ITS-X (10 mg per liter insulin, 5.5 mg per liter transferrin, 0.0067 mg per liter sodium selenite and 2 mg per liter ethanolamine), 8 nM β-estradiol, 200 ng/ml progesterone and 25 μM *N*-acetyl-L-cysteine), as described. Experiments performed in E3.5 ROSA26^DHB/H2B^ embryos with the addition of exogenous FGF4 (1μg/mL) were performed in uncoated 18-wells chambered coverslips (Ibidi) to prevent attachment. To quantify the translocation levels of the CDK sensor during development, two independent datasets with embryos at different stages of development were collected (Figure 2C, N=54; Figure S3C, N=46).

### Immunofluorescence

Mouse and human ESC^DHB/H2B^ were fixed in 2% paraformaldehyde (Electron Microscopy Sciences) for 10 minutes at room temperature and permeabilized for 10 minutes using the following permeabilization buffer (100mM Tris-HCl pH 7.4, 50mM EDTA pH 8.0, 0.5% Triton X- 100). Embryos were fixed in 4% paraformaldehyde for 10 min, permeabilized for 30 minutes in 0.3% Triton X-100 and 0.1 M glycine in 1× phosphate-buffered saline (PBS) and blocked for 1 hour with 1% bovine serum albumin (BSA) and 0.1% Tween in 1× PBS. ESC and embryos were incubated overnight with primary antibodies, washed in 0.1% Tween in 1× PBS, and incubated with the secondary antibody accordingly for one hour at room temperature. For EdU staining, embryos were incubated with 10μM EdU (Click Chemistry Tools) for 30 minutes before fixation. EdU incorporation was visualized using Alexa Fluor 647-azide (Click Chemistry Tools) Click-IT labeling chemistry and Copper protectant to preserve the fluorescence signal from the CDK sensor. Images were acquired using the Nikon SoRa spinning disk confocal microscope equipped with 20x plan-apochromat, 40x plan-apochromat λD 40x OFN25 DIC N2 Air and 60x apochromat TIRF Oil DIC N2 objective lenses (N.A. 0.75, 0.95 and 1.49, respectively) or a Zeiss LSM880 confocal microscope equipped with 20x plan-apochromat and 40x plan-apochromat Oil (N.A. 0.8 and 1.3, respectively) and stage top incubators to maintain temperature, humidity and CO2 concentration (Tokai Hit STX and Okolab Bold Line, respectively). For time-lapse microscopy experiments with embryos, images were acquired every 20 minutes. Image processing was done using Nikon’s NIS-Elements software (v.5.4.1) and images were denoised using the denoise.ai module.

### Western blot

Cells were trypsinized and lysed in 50 mM Tris pH 8, 8 M Urea (Sigma) and 1% Chaps (Millipore) followed by 30 minutes of shaking at 4°C. A total of 20 μg of extracts were run on 4%-12% NuPage Bis-Tris Gel (Invitrogen) and transferred onto Nitrocellulose Blotting Membrane (GE Healthcare). Transferred membranes were incubated with the following primary antibodies overnight at 4°C. The next day the membranes were incubated with HRP-conjugated secondary antibodies Goat anti-Rabbit IgG (H+L) (1:5000; Thermo Fisher Scientific, Cat# 31466) or Goat anti-Mouse IgG (H+L) (1:5000; Thermo Fisher Scientific, Cat# 31431) for 1 hour at room temperature. Membranes were developed using SuperSignal West Pico PLUS or SuperSignal West Femto Maximum sensitivity (Thermo Scientific). For westerns, two independent experiments were performed but only one representative is shown in the figures.

### High throughput imaging (HTI)

A total of 10,000-20,000 human or mouse ESC^DHB/H2B^ (depending on the experiment and on the specific ESC line) were plated on gelatinized μCLEAR bottom 96-well plates (Greiner Bio-One, 655087). ESC were treated with CDK1/2i (different concentrations in the range from 1–30μM depending on the experiment), CDK2i (PF4091 (2 μM –20μM) or CVT313 (200nM–30μM)), CDK1i (RO-3306, 200nM–10μM), Palbociclib (0.2μM–5μM), MYCi (25μM–200μM), mTORi (75nM–500nM) and dTAG (500nM) for one to two hours depending on the experiment before fixation with 4% PFA in PBS for 10 minutes at room temperature. Images were automatically acquired using a CellVoyager CV7000 high throughput spinning disk confocal microscope (Yokogawa, Japan). Each condition was always performed in triplicate wells and at least 9 different fields of view (FOV) were acquired per well. All experiments were performed at least three times with two independent ESC, TSC or hESC clonal lines. Image analysis was performed using the Signals Image Artist system (Revvity Signals). Nuclei were segmented based on H2B- mRuby3. Various measurements were calculated over the nuclear masks, including mean fluorescence intensities from nuclei and external ring around the nuclei. Cell-level data was exported as text files, then analyzed and plotted using R version 4.3.1. When analyzing HTI data, we considered statistically significant those samples that when compared showed an unpaired two-tail t-test with a p-value of at least 0.05 or lower and an averaged Log2 fold change difference of 0.2. For all HTI experiments, three independent experiments including at least two independent clonal cell lines were performed but only one representative experiment is shown in the figures.

### RNA extraction and qPCR

Isolation of total RNA was performed by using the Isolate II RNA Mini Kit (Bioline) followed by cDNA synthesis (SensiFAST cDNA Synthesis Kit, Bioline). Quantitative real time PCR was performed with PowerUp SYBR Master mix in a QuantStudio 6 Pro system. When analyzing quantitative real time PCR data, we considered statistically significant those samples that when compared showed an averaged of two-fold difference in overall gene expression and an unpaired two-tail t-test with a p-value of at least 0.05 or lower. Three independent experiments including at least two independent clonal cell lines were performed but only one is shown in the figures.

### Master Mold Fabrication

The master mold was made from SU-8 photoresist patterned on silicon using standard photolithographic techniques. Briefly, a 4” silicon wafer was dehydrated at 200°C for 20 minutes. To pattern the features, SU-8 2050 (Kayaku Advanced Materials) was spin-coated (Laurell Technologies) on the wafer for a desired height of 37 μm. The wafer was baked at 65°C for 1 minute and then at 95°C for 6 minutes before being exposed with collimated near-UV light through a glass-chrome mask (Front Range Photomask) using a contact mask aligner (OAI Instruments) to form arrays of 1 mm diameter pillars. After exposure, the wafer was again baked at 65°C for 1 minute and then at 95°C for 6 minutes. The unexposed areas of resist were then removed by immersing the wafer in a bath of SU-8 developer (Kayaku Advanced Materials) with gentle agitation for 6 minutes, followed by rinses in isopropanol and then ultrapure water. To increase durability, the master mold was hard baked at 200°C for 4 minutes and then cooled to room temperature at a rate of 150°C/hr. The wafer was then silanized by exposure to tridecafluoro-1,1,2,2-tetrahydrooctyl-1-trichlorosilane (UCT Specialties) vapor overnight to facilitate the unmolding steps. We decided to use a mold of SU-8 pillars followed by an additional PDMS molding step to get PDMS pillars rather than patterning SU-8 wells for two reasons: to avoid adhesion issues arising from in thermal expansion coefficient mismatches when large areas of SU-8 are in contact with the silicon, and to ensure that the pillar edges were sharp and well-defined.

### Two-step Soft Lithography for PDMS Stamp Fabrication

Polydimethylsiloxane/PDMS (Sylgard 184, Dow Corning) base and curing agent were mixed in a 10:1 (w:w) ratio in a planetary mixer (Thinky). 45 g of pre-mixed PDMS was then poured onto the master mold in a 100 mm petri dish, degassed, and cured at 80°C for 1 hour on a leveled shelf. The cured PDMS, with an array of wells, was then unmolded and cut. This PDMS layer was treated with oxygen plasma (Plasma Etch) for 45 seconds and then silanized as described above before being used as a mold for the final PDMS stamps.

### Fabricating PDMS Stamps

PDMS base and curing agent were mixed in a 10:1 (w:w) ratio. 90 g and 70 g of pre-mixed PDMS was poured onto the PDMS mold and into an empty 100 mm petri dish respectively. Both dishes were then degassed and cured as previously described. The plain PDMS sheet served as a backing for the PDMS pillar stamps, with the two layers bonded together after exposure to oxygen plasma for 45 seconds to achieve a total PDMS stamp height of 1-1.5 cm. The additional PDMS height provided ease of handling and sufficient force during the stamping process; we formed the stamp in two layers because it was convenient to use standard height petri dishes while curing the PDMS. PDMS stamps were then cut into individual pieces, approximately 6.5 mm x 6.5 mm and allowed to regain their hydrophobicity for a few days before proceeding to the next steps.

### Gastruloid cell seeding protocol

H9 hESC (WiCell) were seeded and differentiated to generate gastruloids as described. Briefly, single-use PDMS stamps, each with an array of 3 x 3 1mm-diameter pillars, with 1 mm diameter pillars (total of 9 per well) were covered with drops of Matrigel diluted in DMEM-F12 and incubated for 30 minutes at 37°C. This suspension was then aspirated off and the stamps were left to dry at room temperature for up to 30 minutes. Matrigel-coated PDMS stamps were then brought into contact with 8-well Ibidi culture slides for 2 minutes. The printed surface was then washed with DMEM-F12, and wells were ready for cell seeding. Exponentially growing H9 cells at 60-80% confluency were collected by Accutase (Gibco, #A11105-01), washed with 10ml of DMEM-F12 and centrifuged at 1000rpm for 5 minutes. Cells were resuspended in mTeSR Plus containing 10μM Rho-associated kinase inhibitor (ROCKi, Selleckchem, #S1049) and seeded in a single eight-well chamber slide with a total number of 150,000-200,000 cells (Ibidi, #80826). Cells were incubated for six hours at 37°C and 5% CO2, washed gently three times with warm DMEM-F12 and refed with mTeSR without ROCKi for 2 hours followed with mTeSR containing 50ng/ml BMP4 (R&D systems, #314-BP-010) for 42-48 hours. At least two independent experiments with multiple gastruloids per experiment were performed but only one representative is shown in the figures.

### Time-lapse microscopy and single-cell tracking

For all experiments the cells were plated in 96-well optical-grade plastic dishes (Ibidi) at 5000-10,000 cells per well with 300uL media per well. Images were acquired every 12 minutes using a Nikon Ti2-E inverted fluorescent microscope with an automated state and environmental chamber which maintained the cells at 37C and supplied with 5% CO2. Images were acquired in RFP and YFP channels with a 20x 0.45 numerical aperture objective. A total of 6 regions of interest (ROI) per well and a total of 48 wells were imaged for a period of 72 hrs. The total light exposure was less than 500ms for each site that was imaged. Downstream image processing, segmentation, quantification of fluorescent signals (H2B-mRuby3 intensity and CDK-mClover3 activity), and single-cell tracking was performed using custom-written MATLAB scripts as previously described (https://github.com/scappell/Cell_tracking).

### Analysis of single-cell tracking data and single-cell fates

Single cell traces were aligned to the start of the track for individual cells. CDK activity and nuclear area were quantified as previously described (Cappell et al, 2016). Combined single cell traces were aligned to the time of mitosis for each individual cells and cell fates were quantified as followed: CDK increasing (CDK inc) cells are defined as CDK activity > −0.74 within 4hrs of mitosis. CDK delayed cells are defined as CDK < −0.74 for the first 4hrs following mitosis and then subsequent CDK activity > −0.74. G1 exit is defined as CDK < −0.74 for the duration of the observation period. G2 exit is defined as a CDK > −0.74 after mitosis, followed by CDK < −0.74 for the remainder of the observation period. Endoreduplication is defined as exit from G2 (CDK < − 0.74) followed by re-accumulation of CDK2 activity, without mitosis.

### Image analysis for mouse embryos

Image analysis and segmentation was performed using custom scripts written in Python (v. 3.10). Deep learning segmentation models were trained and deployed using the NIH HPC Biowulf cluster (http://hpc.nih.gov). H2B-mRuby3 labeled embryonic nuclei were segmented using a custom trained Cellpose model (Pachitariu and Stringer, 2022), creating individual nuclear ROIs. To create the cytoplasmic ROI, each nuclear label was expanded by 2 pixels and 12 pixels. The 2-pixel expansion was subtracted from the 12-pixel expansion, resulting in a cytoplasmic ROI, matched to a parental nuclear ROI. Both the nuclear and cytoplasmic ROIs were used to quantify fluorescent intensities across the other imaged markers (DHB-mClover3, CDX2 or SOX2). DHB- mClover3 cytoplasmic and nuclear intensities were used to calculate a cytoplasmic/nuclear ratio. Embryonic cells were classified as either CDX2 or SOX2 positive or negative by setting a fluorescent intensity threshold based on the distribution of nuclear marker intensities. All cells were linked to their parent embryo, allowing a cell count per embryo.

### Image analysis for human gastruloids

Gastruloid image analysis and segmentation was performed using custom scripts written in Python (v. 3.10). Deep learning segmentation models were trained and deployed using the NIH HPC Biowulf cluster (http://hpc.nih.gov). H2B-mRuby3 labeled gastruloid nuclei were segmented using a custom trained Cellpose model, creating individual nuclear ROIs. Cytoplasmic ROIs were generated as described for embryos, by extending the nuclear labels by 12-pixels and subtracting the nuclear + 2-pixel label. Cytoplasmic and nuclear ROIs were used to quantify marker intensities as described above. The entire gastruloid was also segmented in order to identify the gastruloid centroid. The position of every nucleus relative to the gastruloid centroid was calculated using a Euclidean distance transform. Nuclear distances were either binned into 5 concentric rings every 100 um away from the centroid (A = 0 – 100 μm, B = 100 – 200 μm, C = 200 – 300 μm, D = 300 – 400 μm, E >= 400 μm) or nuclear distances were normalized into radial distances (0 – 1) by calculating nuclear distance from centroid / maximum gastruloid radial distance. Cells were classified into CDX2, SOX2, TFAP2C, and BRA positive or negative by setting a fluorescent intensity threshold based on the distribution of nuclear marker intensities.

### Lead contact

All requests for additional information and resources should be directed to Sergio Ruiz Macias.

### Materials availability

Reagents that support the findings of this study are available upon reasonable request from the lead contact.

## REFERENCES

Aleem, E., Kiyokawa, H. and Kaldis, P. (2005) Cdc2-cyclin E complexes regulate the G1/S phase transition. Nat. Cell Biol. 7, 831–836.

Bedzhov, I., Leung, C.Y., Bialecka, M. and Zernicka-Goetz, M. (2014) In vitro culture of mouse blastocysts beyond the implantation stages. Nat. Protoc. 9, 2732–2739.

Boroviak, T., Loos, R., Lombard, P., Okahara, J., Behr, R., Sasaki, E., Nichols, J., Smith, A. and Bertone, P. (2015) Lineage-specific profiling delineates the emergence and progression of naive pluripotency in mammalian embryogenesis. Dev. Cell 35, 366–382.

Bulut-Karslioglu, A., Biechele, S., Jin, H., Macrae, T.A., Hejna, M., Gertsenstein, M., Song, J.S. and Ramalho-Santos M. (2016) Inhibition of mTOR induces a paused pluripotent state. Nature 540, 119–123.

Cappell, S.D., Chung, M., Jaimovich, A., Spencer, S.L. and Meyer, T. (2016) Irreversible APC(Cdh1) Inactivation Underlies the Point of No Return for Cell-Cycle Entry. Cell 166, 167–180.

Chazaud, C. and Yamanaka, Y. (2016) Lineage specification in the mouse preimplantation embryo. Development, 143, 1063–1074.

Ciemerych, M.A. and Sicinski, P. (2005) Cell cycle in mouse development. Oncogene 24, 2877–98.

Christodoulou, N., Weberling, A., Strathdee, D., Anderson, K.I., Timpson, P. and Zernicka-Goetz, M. (2019) Morphogenesis of extra-embryonic tissues directs the remodeling of the mouse embryo at implantation. Nat. Commun. 10: 3557.

Copp, A.J. (1978) Interaction between inner cell mass and trophectoderm of the mouse blastocyst. I. A study of cellular proliferation. J. Embryol. Exp. Morphol. 48, 109–125.

Fischer, M., Schade, A.E., Branigan, T.B., Müller, G.A. and DeCaprio, J.A. (2022) Coordinating gene expression during the cell cycle. Trends Biochem. Sci. 47, 1009–1022.

Iwamori, N., Naito, K., Sugiura, K. and Tojo, H. (2002) Preimplantation-embryo-specific cell cycle regulation is attributed to the low expression level of retinoblastoma protein. FEBS Lett. 526, 119–23.

Kalkan, T., Olova, N., Roode, M., Mulas, C., Lee, H.J., Nett, I., Marks, H., Walker, R., Stunnenberg, H.G., Lilley, K.S., Nichols, J., Reik, W., Bertone, P. and Smith, A. (2017) Tracking the embryonic stem cell transition from ground state pluripotency. Development 144, 1221–1234.

Malumbres, M. and Barbacid, M. (2009) Cell cycle, CDKs and cancer: a changing paradigm. Nat. Rev. Cancer 9, 153–66.

Massague, J. G1 cell-cycle control and cancer. (2004) Nature 432, 298–306.

Michowski, W., Chick, J.M., Chu, C., Kolodziejczyk, A., Wang, Y., Suski, J.M., Abraham, B., Anders, L., Day, D., Dunkl, L.M., Li, Cheong Man, M., Zhang, T., Laphanuwat, P., Bacon, N.A., Liu, L., Fassl, A., Sharma, S., Otto, T., Jecrois, E., Han, R., Sweeney, K.E., Marro, S., Wernig, M., Geng, Y., Moses, A., Li, C., Gygi, S.P., Young, R.A. and Sicinski, P. (2020) Cdk1 Controls Global Epigenetic Landscape in Embryonic Stem Cells. Mol. Cell 78, 459–476.

Minn, K.T., Fu, Y.C., He, S., Dietmann, S., George, S.C., Anastasio, M.A., Morris, S.A. and Solnica-Krezel, L. (2020) High-resolution transcriptional and morphogenetic profiling of cells from micropatterned human ESC gastruloid cultures. Elife. 9: e59445.

Nakayama, K. and Nakayama, K. (1998) Cip/Kip cyclin-dependent kinase inhibitors: brakes of the cell cycle engine during development. Bioessays 20, 1020–1029.

Liu, L., Michowski, W., Kolodziejczyk, A. and Sicinski, P. (2019) The cell cycle in stem cell proliferation, pluripotency and differentiation. Nat. Cell Biol. 21, 1060–1067.

Pachitariu, M. and Stringer, C. (2022) Cellpose 2.0: how to train your own model. Nat. Methods. 19, 1634–1641.

Padgett, J. and Santos, S.D.M. (2020) From clocks to dominoes: lessons on cell cycle remodelling from embryonic stem cells. FEBS Lett. doi: 10.1002/1873-3468.13862.

Pauklin, S. and Vallier, L. (2013) The cell-cycle state of stem cells determines cell fate propensity. Cell 155, 135–147.

Pokrass, M.J., Ryan, K.A., Xin, T., Pielstick, B., Timp, W., Greco, V. and Regot, S. (2020) Cell-Cycle-Dependent ERK Signaling Dynamics Direct Fate Specification in the Mammalian Preimplantation Embryo. Dev. Cell 55, 328–340.

Rossant, J. (2018) Genetic Control of Early Cell Lineages in the Mammalian Embryo. Annu. Rev. Genet. 52, 185–201.

Ruiz, S., Panopoulos, A.D., Herrerías, A., Bissig, K.D., Lutz, M., Berggren, W.T., Verma, I.M. and Izpisua Belmonte, J.C. (2011) A high proliferation rate is required for cell reprogramming and maintenance of human embryonic stem cell identity. Curr. Biol. 2, 45–52.

Shahbazi, M.N. and Zernicka-Goetz, M. (2018) Deconstructing and reconstructing the mouse and human early embryo. Nat. Cell Biol. 20, 878–887.

Savatier, P., Lapillonne, H., van Grunsven, L.A., Rudkin, B.B. and Samarut, J. (1996) Withdrawal of differentiation inhibitory activity/leukemia inhibitory factor up-regulates D-type cyclins and cyclin-dependent kinase inhibitors in mouse embryonic stem cells. Oncogene 12, 309–22.

Scognamiglio, R., Cabezas-Wallscheid, N., Their, M.C., Altamura, S., Reyes, A., Prendergast, Á.M., Baumgärtner, D., Carnevalli, L.S., Atzberger, A., Haas, S., von Paleske, L., Boroviak, T., Wörsdörfer, P., Essers, M.A., Kloz, U., Eisenman, R.N., Edenhofer, F., Bertone, P., Huber, W., van der Hoeven, F., Smith, A. and Trumpp A. (2016) Myc Depletion Induces a Pluripotent Dormant State Mimicking Diapause. Cell 164, 668–80.

Spencer, S.L., Cappell, S.D., Tsai, F.C., Overton, K.W., Wang, C.L. and Meyer, T. (2013) The proliferation-quiescence decision is controlled by a bifurcation in CDK2 activity at mitotic exit. Cell 155, 369–83.

Stead, E., White, J., Faast, R., Conn, S., Goldstone, S., Rathjen, J., Dhingra, U., Rathjen, P., Walker, D. and Dalton, S. (2002) Pluripotent cell division cycles are driven by ectopic Cdk2, cyclin A/E and E2F activities. Oncogene 21, 8320–8333.

Stewart, C.L. and Cullinan, E.B. (1997) Preimplantation development of the mammalian embryo and its regulation by growth factors. Dev. Genet. 21, 91–101.

Simon, C.S., Rahman, S., Raina, D., Schröter, C. and Hadjantonakis, A.K. (2020) Live Visualization of ERK Activity in the Mouse Blastocyst Reveals Lineage-Specific Signaling Dynamics. Dev. Cell 55, 341–353.

Tanaka, S., Kunath, T., Hadjantonakis, A. K., Nagy, A. and Rossant, J. (1998) Promotion of trophoblast stem cell proliferation by FGF4. Science 282, 2072–2075.

Ullah, Z., Kohn, M.J., Yagi, R., Vassilev, L.T. and DePamphilis, M.L. (2008) Differentiation of trophoblast stem cells into giant cells is triggered by p57/Kip2 inhibition of CDK1 activity. Genes Dev. 22, 3024–36.

Wang, X.Q., Lo, C.M., Chen, L., Ngan, E.S., Xu, A. and Poon, R.Y. (2017) CDK1-PDK1-PI3K/Akt signaling pathway regulates embryonic and induced pluripotency. Cell Death Differ. 24, 38–48.

Warmflash, A., Sorre, B., Etoc, F., Siggia, E.D. and Brivanlou, A.H. (2014) A method to recapitulate early embryonic spatial patterning in human embryonic stem cells. Nat. Methods. 11, 847–854.

White, J. and Dalton, S. (2005) Cell cycle control of embryonic stem cells. Stem Cell Rev. 1, 131–138.

White, J., Stead, E., Faast, R., Conn, S., Cartwright, P. and Dalton, S. (2005) Developmental activation of the Rb-E2F pathway and establishment of cell cycle-regulated cyclin-dependent kinase activity during embryonic stem cell differentiation. Mol. Biol. Cell 16, 2018–27.

Williams, R.L., Hilton, D.J., Pease, S., Willson, T.A., Stewart, C.L., Gearing, D.P., Wagner, E.F., Metcalf, D., Nicola, N.A. and Gough, N.M. (1988) Myeloid leukemia inhibitory factor maintains the developmental potential of embryonic stem cells. Nature 336, 684–687.

Yao, C., Zhang, W. and Shuai, L. (2019) The first cell fate decision in pre-implantation mouse embryos. Cell Regen. 8, 51–57.

