## Supplementary Figures 1-6 for "Lineage-specific CDK activity dynamics characterize early mammalian development"

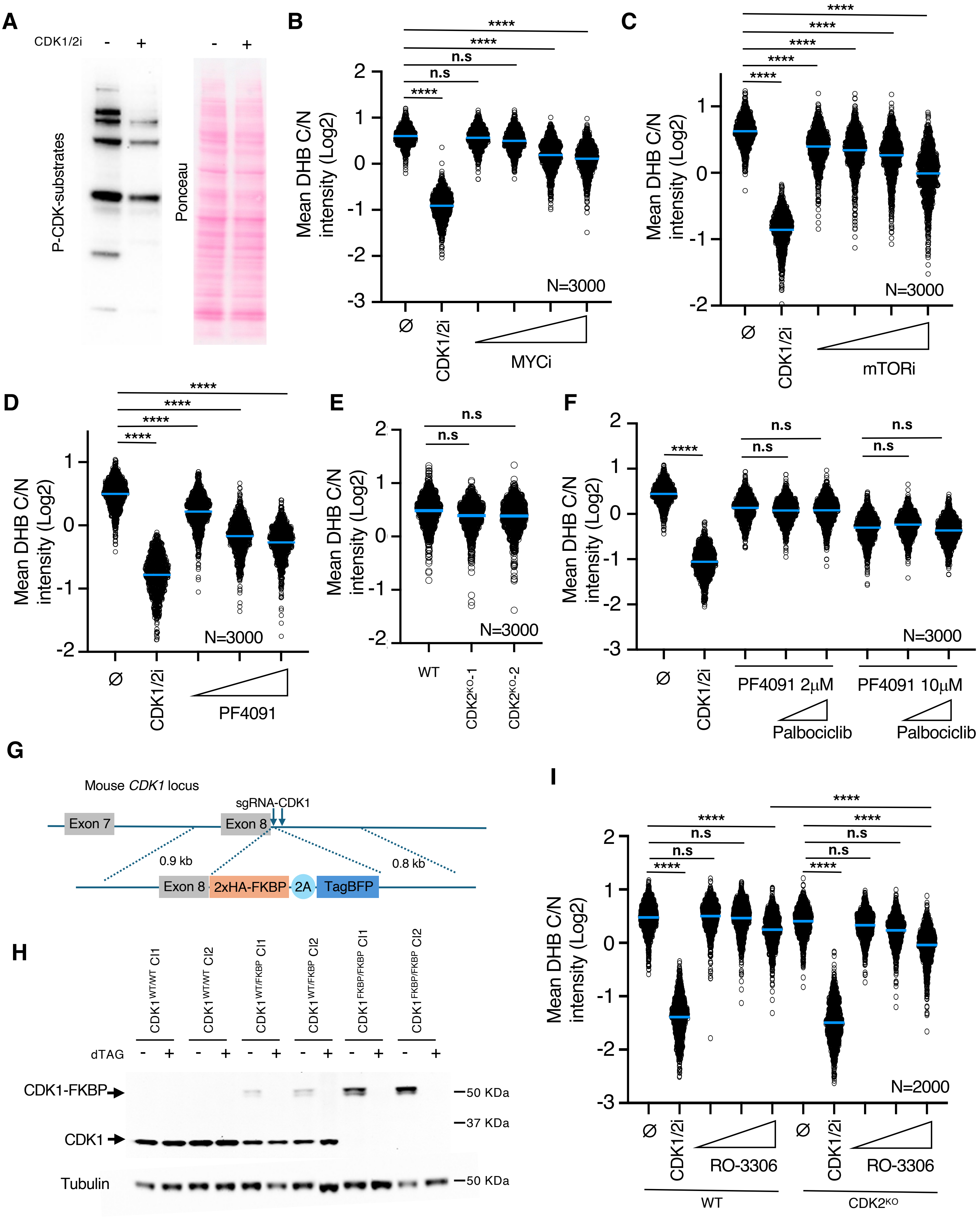


Saykali et al, Supplementary Figure 1

**Figure S1: The CDK-KTR sensor responds to changes in CDK2 and CDK1 activity.**

(A) Western blot analysis to detect phospho (P)-CDK substrates using lysates from CDK1/2i-treated (30μM) $ESC^{DHB/H2B}$ cultures for one hour.

(B and C) High-throughput imaging (HTI) quantification of C/N mean intensity in untreated (⌀) and MYCi (25μM, 50μM, 100μM and 200μM) (A) or mTORi (75nM, 125nM, 250nM and 500 nM) (B)-treated $ESC^{DHB/H2B}$ cultures with increasing concentration of the inhibitor for 24 hours. Center lines indicate mean values. N=3000 cells; p-values are shown from two-tailed unpaired *t*-tests. **** p<0.0001; n.s.: non-significant.

(D) HTI quantification of C/N mean intensity in untreated (⌀) and PF4091-treated $ESC^{DHB/H2B}$ cultures with increasing concentration of the inhibitor (2μM, 10μM, and 20μM) for one hour. Center lines indicate mean values. N=3000 cells; p-values are shown from two-tailed unpaired *t*-tests. **** p<0.0001; n.s.: non-significant.

(E) HTI quantification of C/N mean intensity in wild-type or CDK2-deficient $ESC^{DHB/H2B}$ cultures for one hour. Center lines indicate mean values. N=3000 cells; p-values are shown from two-tailed unpaired *t*-tests. **** p<0.0001.

(F) HTI quantification of C/N mean intensity in untreated (⌀) and PF4091-treated (10μM) $ESC^{DHB/H2B}$ cultures combined or not with Palbociclib at different concentrations (1μM and 5μM) for one hour. Center lines indicate mean values. N=3000 cells; p-values are shown from two-tailed unpaired *t*-tests. **** p<0.0001; n.s.: non-significant.

(G) Schematic representation of the targeting event on the endogenous CDK1 locus using the depicted recombination construct. Two different sgRNAs were used to induce the targeting.

(H) Western blot analysis to evaluate CDK1 degradation upon 500nM dTAG treatment in wild type, heterozygous and homozygous $ESC^{DHB/H2B}$ clonal lines.

(I) HTI quantification of C/N mean intensity in untreated (⌀) and RO-3306-treated wild-type or CDK2-deficient $ESC^{DHB/H2B}$ cultures at different concentrations (5μM, 10μM, and 30μM) for one hour. $ESC^{DHB/H2B}$ treated with CDK1/2i (30μM) were used as control. Center lines indicate mean values. N=2000 cells; p-values are shown from two-tailed unpaired *t*-tests. **** p<0.0001; n.s.: non-significant.

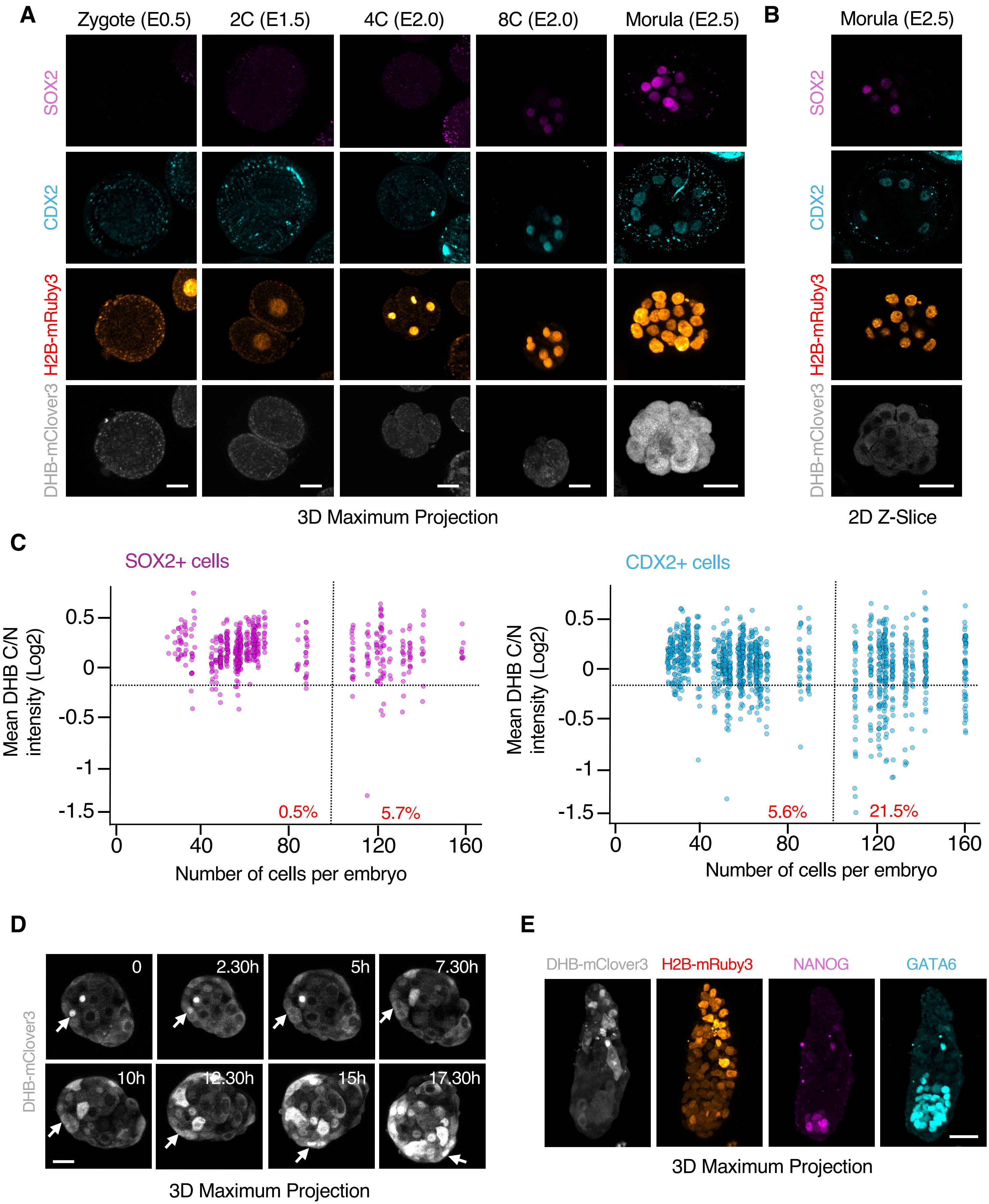


Saykali et al, Supplementary Figure 2

**Figure S2: Cells from TE lower CDK activity prior to implantation**

(A and B) Confocal images from hemizygous ROSA26$^{DHB/H2B}$ embryos during pre-implantation development (from zygote to morula). Scale bar, 30μm (A) and 20μm (B).

(C) Plot showing a quantification of C/N mean intensity in individual cells (each represented by a dot) obtained from hemizygous ROSA26$^{DHB/H2B}$ embryos during pre-implantation development. The collection of dots from the same column belongs to embryos with same cell count. Embryos were staged based on the number of cells and cells classified as SOX2 (left panel) or CDX2 (right panel) expressing cells in each embryo. In red is the percentage of cells below the defined arbitrary threshold (-0.25) shown for embryos containing 0-100 cells (left) and embryos containing above 100 cells (right). Data from this plot was derived from an independent set of embryos from those shown in Figure 2C. N=46 embryos.

(D) Confocal images from a representative hemizygous ROSA26$^{DHB/H2B}$ embryo isolated at E3.5 where an event of endoreplication could be detected (arrows). Scale bar, 30μm.

(E) Confocal images from a representative hemizygous ROSA26$^{DHB/H2B}$ embryos isolated at E4.5. Scale bar, 30μm.

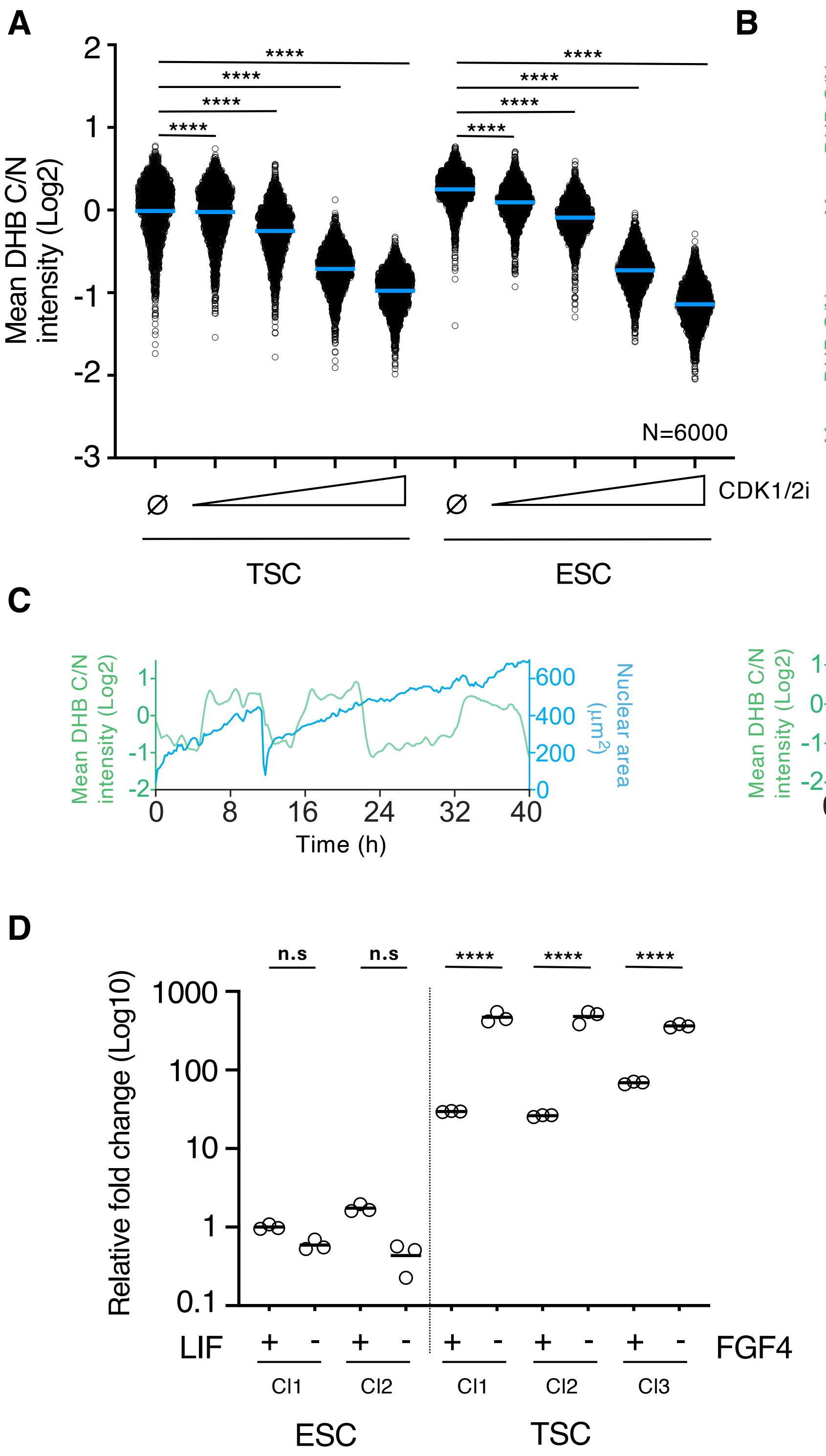


Saykali et al, Supplementary Figure 3

**Figure S3: FGF4 withdrawal leads to endoreplication and p57$^{KIP2}$ upregulation in TSC**

(A) High-throughput imaging (HTI) quantification of C/N mean intensity in untreated (⌀) or CDK1/2i-treated TSC$^{DHB/H2B}$ and ESC$^{DHB/H2B}$ cultures with increasing concentration of the inhibitor (200nM, 1μM, 10μM and 30μM) for one hour. Center lines indicate mean values. N=6000 cells; p-values are shown from two-tailed unpaired *t*-tests. **** p<0.0001; n.s.: non-significant.

(B and C) Single cell CDK activity traces of two representative examples of proliferating (B) or 72 hours FGF4-deprived (C) TSC$^{DHB/H2B}$. In (C), one cell undergoing G1 exit (left track) or endoreplication (right track) are shown. Nuclear size was also quantified over time.

(D) Relative fold change (log10) expression of *p57$^{KIP2}$* in ESC$^{DHB/H2B}$ and TSC$^{DHB/H2B}$ growing under self-renewal conditions or deprived of LIF or FGF4, respectively. Reactions were performed in triplicate using at least two different clonal lines. p-values are shown from two-tailed unpaired *t*-tests. **** p<0.0001; n.s.: non-significant.

(E) Confocal images of TSC$^{DHB/H2B}$ growing under self-renewal conditions or FGF4-deprived for 72 hours. Scale bar, 25μm.

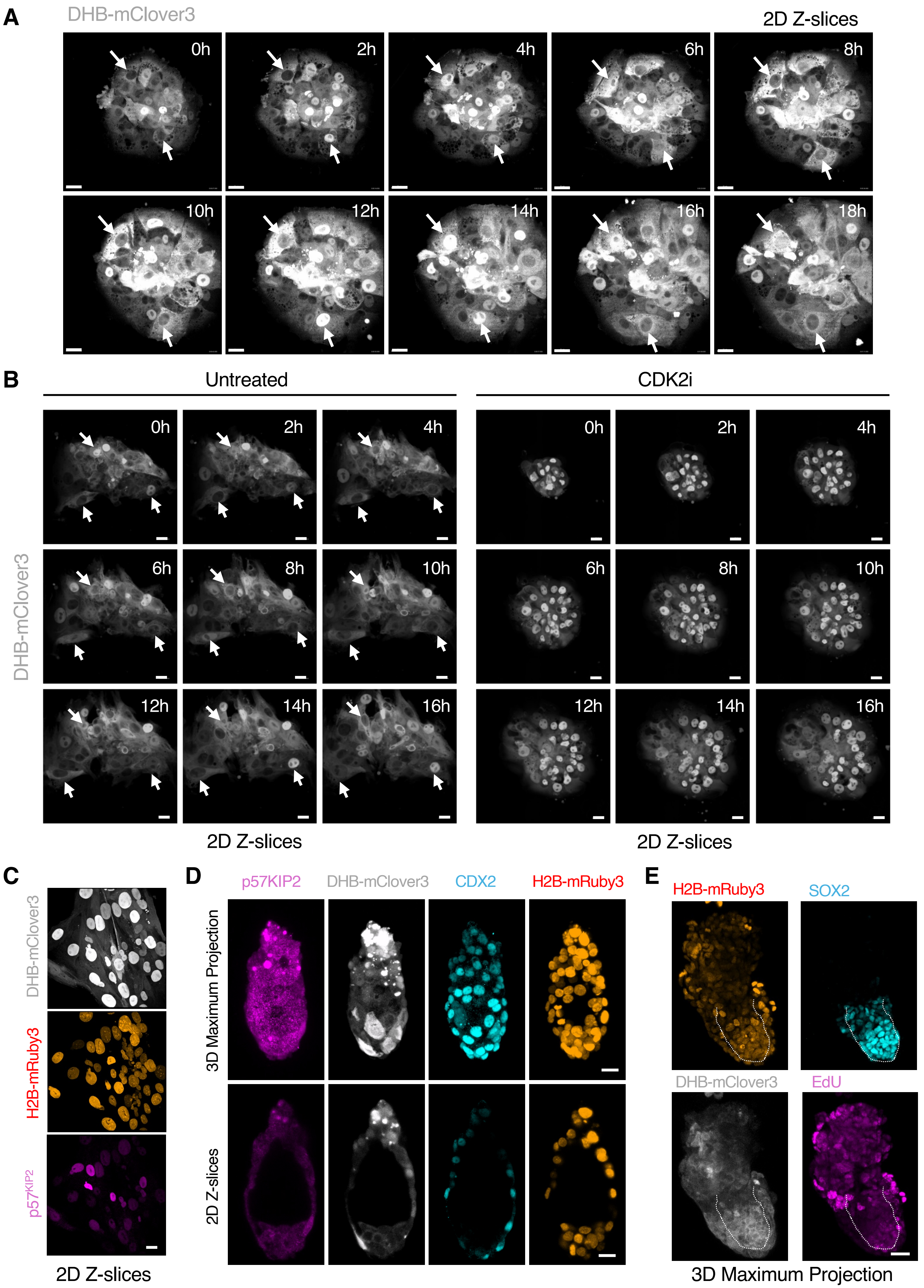


Saykali et al, Supplementary Figure 4

**Figure S4: Endoreplication in TGC from ROSA26$^{DHB/H2B}$ embryos cultured beyond implantation stages.**

(A) Time-lapse microscopy imaging performed in attached hemizygous ROSA26$^{DHB/H2B}$ embryos 48 hours after isolation (E3.5). One representative embryo is shown. Arrows indicate several examples of cells undergoing endoreplication. Scale bar, 30μm.

(B) Time-lapse microscopy imaging performed in untreated or 10μM PF4091-treated attached hemizygous ROSA26$^{DHB/H2B}$ embryos for 36 hours following isolation at E3.5. Representative embryos are shown. Arrows indicate several examples of cells undergoing endoreplication. Scale bar, 50μm.

(C) Confocal images of a representative attached hemizygous ROSA26$^{DHB/H2B}$ embryos 48 hours after isolation at E3.5. Scale bar, 30μm.

(D) Confocal images of representative E4.5 ROSA26$^{DHB/H2B}$ embryos. Scale bar, 20μm.

(E) Confocal images of a representative hemizygous ROSA26$^{DHB/H2B}$ embryo cultured four days after isolation at E3.5. Scale bar, 30μm.

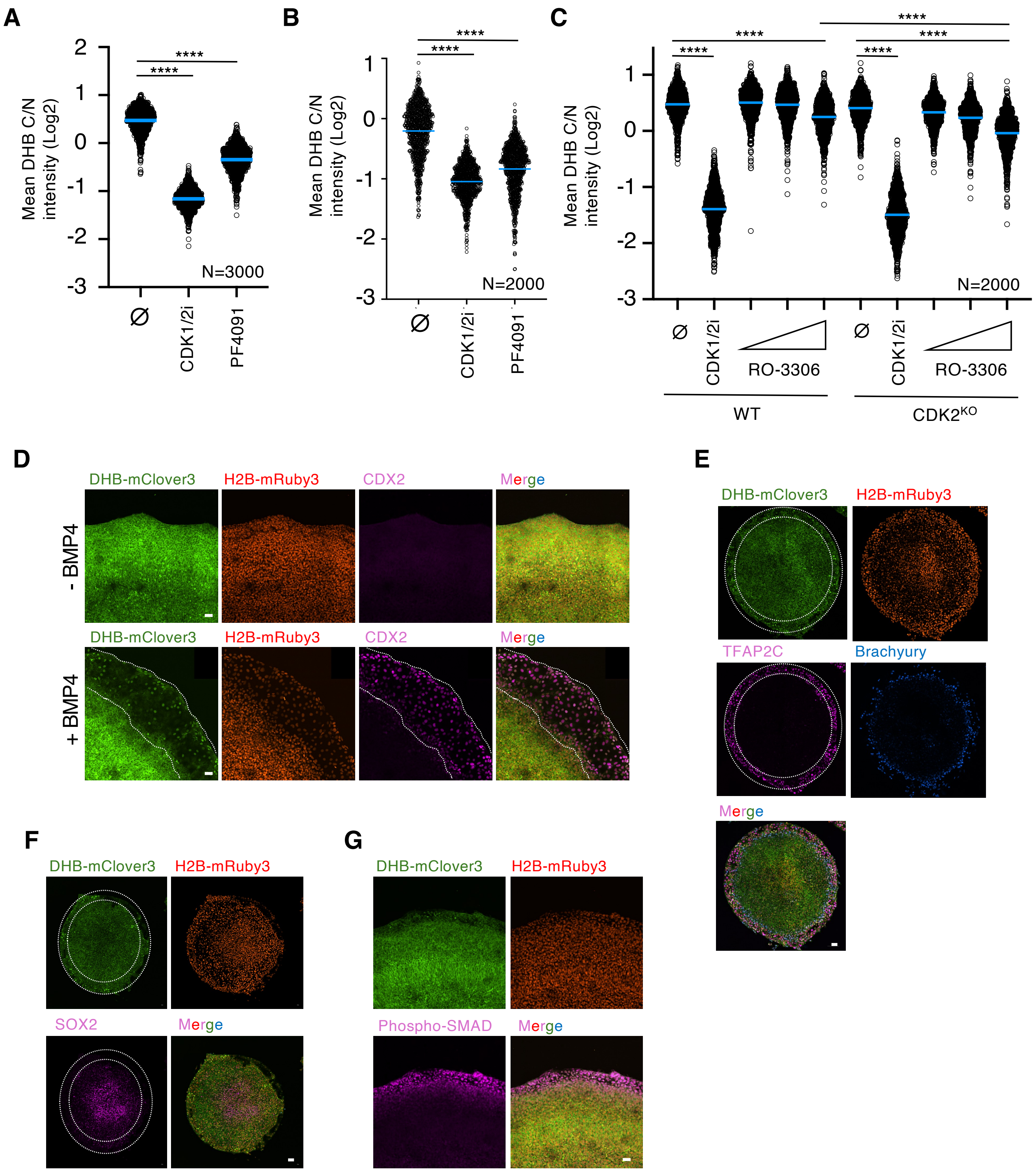


Saykali et al, Supplementary Figure 5

**Figure S5: BMP4-induced gastruloids from hESC as a model to analyze dynamics of the CDK sensor.**

(A, B) High-throughput imaging (HTI) quantification of C/N mean intensity in untreated (⌀) or 20μM PF4091-treated primed (A) or naïve (B) $hESC^{DHB/H2B}$ cultures for one hour. CDK1/2i-treated $hESC^{DHB/H2B}$ (30μM) cultures were used as control. Center lines indicate mean values. N=3000 cells (A); N=2000 cells (B). p-values are shown from two-tailed unpaired *t*-tests. **** p<0.0001.

(C) HTI quantification of C/N mean intensity in untreated (⌀) and RO-3306-treated wild-type or CDK2-deficient $hESC^{DHB/H2B}$ cultures at different concentrations (5μM, 10μM and 30μM) for one hour. $hESC^{DHB/H2B}$ treated with CDK1/2i (30μM) were used as control. Center lines indicate mean values. N=2000 cells; p-values are shown from two-tailed unpaired *t*-tests. **** p<0.0001. Only significant comparisons are shown.

(D-G) Confocal images of representative gastruloids uninduced or BMP4-induced for seventy-two hours. Dashed lines indicate the outer ring of TE-like cells identified by CDX2 expression. Scale bar, 100μm (E, F), 50μm (D, G).

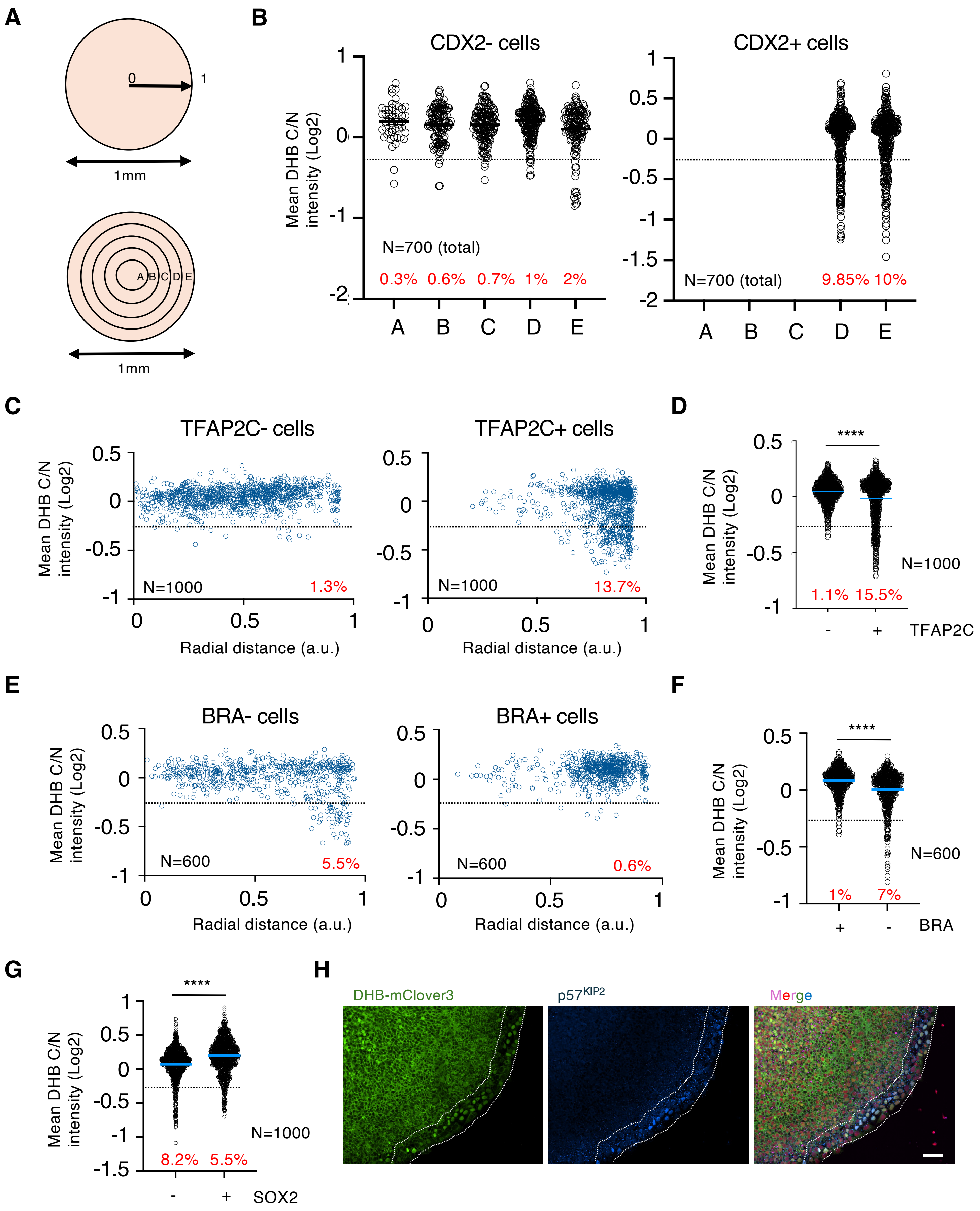


Saykali et al, Supplementary Figure 6

**Figure S6: Human TE-like cells accumulate the CDK sensor in the nucleus upon differentiation.**

(A) Schematic representation of the circular surface of the gastruloid using radial distance (upper panel) or dividing the circle in concentrical rings (lower panel).

(B) Plot showing quantification of C/N mean intensity in a total number of 700 random CDX2+ and CDX2- individual cells distributed accordingly to their position from the center of the gastruloid. In red is the percentage of cells per ring below the arbitrary threshold (<-0.25). Data was obtained by combining multiple gastruloids from the same experiment (N=5 gastruloids).

(C) Plot showing a quantification of C/N mean intensity in 1000 random TFAP2C+ or TFAP2C- cells distributed accordingly to their radial distance from the center of the gastruloid. In red is the percentage of cells below the arbitrary threshold (<-0.25). Data was obtained by combining multiple gastruloids from the same experiment (N=11 gastruloids).

(D) Plot showing a quantification of C/N mean intensity in 1000 random TFAP2C+ or TFAP2C- cells from human gastruloids. In red is the percentage of cells below the arbitrary threshold (<-0.25). p values are shown from two-tailed unpaired *t*-tests. **** p<0.0001. Data was obtained by combining multiple gastruloids from the same experiment (N=11 gastruloids).

(E) Plot showing a quantification of C/N mean intensity in 600 random Brachyury (BRA)+ or BRA- cells distributed accordingly to their radial distance from the center of the gastruloid. In red is the percentage of cells below the arbitrary threshold (<-0.25). Data was obtained by combining multiple gastruloids from the same experiment (N=3 gastruloids).

(F) Plot showing a quantification of C/N mean intensity in 1000 random Brachyury (BRA)+ or BRA- cells from human gastruloids. In red is the percentage of cells below the arbitrary threshold (<-0.25). p values are shown from two-tailed unpaired *t*-tests. **** p<0.0001. Data was obtained by combining multiple gastruloids from the same experiment (N=3 gastruloids).

(G) Plot showing a quantification of C/N mean intensity in 1000 random SOX2+ or SOX2- cells from human gastruloids. In red is the percentage of cells below the arbitrary threshold (<-0.25). p values are shown from two-tailed unpaired *t*-tests. **** p<0.0001. Data was obtained from combining multiple gastruloids from the same experiment (N=11 gastruloids).

(H) Confocal images of representative BMP4-induced gastruloids for 72 hours. Dashed lines indicate the outer ring of TE-like. Note the specific expression of $p57^{KIP2}$ in cells with nuclear accumulation of the CDK sensor. Scale bar, 50μm.
